# Regulation of the cell surface expression of classical and non-classical MHC proteins by the human cytomegalovirus UL40 and rhesus cytomegalovirus rh67 proteins

**DOI:** 10.1101/2022.07.22.501209

**Authors:** Simon Brackenridge, Persephone Borrow, Andrew McMichael

## Abstract

The signal sequences of the human cytomegalovirus (CMV) UL40 protein, and its rhesus CMV counterpart, Rh67, contain a peptide (VMAPRT[L/V][F/I/L/V]L, VL9) that can be presented by Major Histocompatibility Complex (MHC) antigen E. The CMV VL9 peptides replace VL9 peptides derived from classical MHC (Ia) signal sequences, which are lost when CMV disrupts MHC Ia expression, as well as antigen processing and presentation. This allows infected cells to maintain surface expression of MHC-E and escape killing by NK cells expressing the inhibitory NKG2A/CD94 receptor. We demonstrate that processing of the Rh67 VL9 peptide mirrors that of UL40, despite the lack of sequence conservation elsewhere in the two proteins. As previously shown for UL40, up-regulation of MHC-E expression by Rh67 only requires its signal sequence, with sequences upstream of VL9 critical for conferring independence from TAP, the Transporter Associated with Antigen Processing. Additionally, we show that processing of both VL9 peptides depends on cleavage of the signal sequences by the host protease Signal Peptide Peptidase. Notably, our results also reveal that the mature UL40 and Rh67 proteins contribute to CMV immune evasion by down-regulating surface expression of MHC Ia. Unexpectedly, while the Rh67 VL9 peptide is resistant to the effects of the Rh67 protein, the UL40 protein is able to counteract up-regulation of MHC-E expression mediated by its own VL9 peptide. This suggests differences in the mechanisms by which the two VL9 peptides up-regulate MHC-E, which may have implications for translating a RhCMV-vectored SIV vaccine to HIV-1 using HCMV as a vector.

**IMPORTANCE:** Cell surface MHC-E expression, which requires a peptide (VMAPRT[L/V][F/I/L/V]L, “VL9”) from the signal sequences of other MHC class I proteins, prevents cells from being killed by CD94/NKG2A-expressing Natural Killer cells. In cells infected with human CMV, the endogenous VL9 peptide is replaced by one from the signal sequence of the HCMV UL40 protein. We show that processing of the VL9 peptide of Rh67, the rhesus CMV equivalent of UL40, mirrors that of UL40, despite a lack of sequence homology between the two proteins. Of note, we also show that the mature UL40 and Rh67 proteins, which have no previously described function, contribute to CMV immune evasion by reducing classical MHC class I surface expression. Importantly, the mature UL40 protein, but not the mature Rh67 protein, can decrease the up-regulation of MHC-E mediated by its signal sequence VL9 peptide, which may have implications for HCMV as a vaccine vector.

## INTRODUCTION

Cell surface expression of the Major Histocompatibility (MHC) E protein, known as HLA-E in humans, depends on a nonamer peptide (VMAPRT[L/V][F/I/L/V]L, VL9) present in the signal sequences of many MHC class Ia (MHC Ia) allotypes (1, 2). Presentation of this peptide is dependent on cleavage by both the endoplasmic reticulum (ER) membrane resident proteases Signal Peptidase (SP) and Signal Peptide Peptidase (SPP) (3, 4) and the cytosolic proteasome (5), import into the ER lumen by the transporter associated with antigen presentation (TAP), as well as tapasin and the peptide loading complex (PLC) (6).

The presence of HLA-E at the cell surface is monitored by NK cells via CD94/NKG2 receptor complexes (7, 8) as a proxy for MHC class I expression. Loss of surface HLA-E results in the loss of signalling through the inhibitory NKG2A/CD94 complex, allowing NK cells to kill HLA-E-deficient cells (9). Cell surface expression of HLA-E is, therefore, often maintained by virally infected cells, in contrast to that of classical MHC class I proteins. For example, evasion of CD8^+^ T cell responses by human CMV (HCMV) involves not only direct targeting of the MHC Ia proteins themselves, including retention in the ER by US10 (10) and targeting for degradation by US2 and US11 (11, 12), but also targeting of accessory factors. US6 blocks peptide transport by TAP (13), while US3 inhibits tapasin (14). Such comprehensive disruption of both MHC Ia expression and components of the PLC limits the availability of the VL9 peptide for HLA-E, resulting in NK cell-mediated killing of HCMV infected cells. However, the signal sequence of the HCMV UL40 protein contains a VL9 peptide, which can be presented by HLA-E in a TAP-independent manner (15, 16), allowing HCMV-infected cells to maintain surface expression of HLA-E and avoid detection by NK cells.

Immune evasion by rhesus CMV (RhCMV) appears to be highly conserved, and equivalents of the HCMV US2, US3, US6, and US11 proteins have been identified that function by conserved mechanisms: Rh182, Rh184, Rh186, and Rh188, respectively (17). RhCMV also appears to evade NK cells in the same way as HCMV, via the Rh67 protein. Despite otherwise appearing unrelated to UL40, Rh67 also contains the VL9 peptide in its signal sequence (18), and we previously showed that the Rh67 VL9 peptide can stabilise HLA-E at the cell surface (19). More recently, it has recently been demonstrated that presentation of the Rh67 VL9 peptide does not require TAP (20), confirming that Rh67 is indeed the RhCMV equivalent of UL40.

A RhCMV-vectored vaccine containing the SIV Gag, Rev/Tat/Nef, Env and Pol sequences elicits an immune response that enables ∼55% of vaccinated rhesus macaques to clear SIV infection early after virus challenge (21, 22). Unusually, the CD8+ T cell responses induced by this vaccine are not restricted by classical MHC Ia proteins. Instead, approximately two-thirds of the CD8^+^ T cells are restricted by MHC class II, with the rest restricted by Mamu-E, the rhesus macaque homologue of HLA-E (19). A full understanding of exactly how protection is mediated by this vaccine remains to be elucidated, but it has recently been shown to depend on the Mamu-E-restricted responses rather than the MHC class II-restricted responses (23, 24).

Successful development of a HCMV-vectored HIV-1 vaccine will likely depend on a detailed understanding of the similarities and differences in the immune evasion strategies employed by RhCMV and HCMV. Indeed, such an understanding may even allow the use of HCMV to be avoided, with the required CMV genes being engineered into a more conventional vaccine vector. To this end, we have undertaken a detailed comparison of how the VL9 peptides from UL40 and Rh67 are processed for presentation by HLA-E. Our results reveal that the similarities between UL40 and Rh67 extend beyond their independence from TAP. Processing of both the UL40 and Rh67 VL9 peptides is dependent on cleavage by SPP, and the up-regulation of HLA-E mediated by Rh67 requires only the signal sequence, as was reported previously for UL40 (16). Furthermore, we show that the mature Rh67 and UL40 proteins also contribute to the evasion of the host immune response by decreasing surface expression of MHC Ia. Unexpectedly, the mature UL40 and Rh67 proteins are also capable of counteracting the up-regulation of HLA-E mediated by the UL40 signal sequence, but not the Rh67 signal sequence. This suggests that the TAP-independence of the UL40 and Rh67 VL9 peptides may be achieved by distinct mechanisms, and that there may be differences in the level of MHC-E expression required to evade NK cells in humans and macaques.

## METHODS

### Expression Plasmids and cloning

The single chain dimer (SCD) of HLA-E*01:01 and β2-microglobulin was as previously described (19), and the HLA-E*01:03 SCD was made by an identical cloning strategy. The coding sequences of HLA-A*02:01 (with its own signal sequence, or the signal sequences of UL40 or Rh67), KIR2DS2*001 (used as the irrelevant control), UL40 (HCMV strain Toledo), and Rh67 (RhCMV strain 68-1) were purchased as synthetic genes from Eurofins with appropriate restriction sites (HindIII and BamHI) and inserted into pEGFP-N1 (Clontech). Constructs to express peptides in the cytosol were created by inserting annealed oligonucleotides comprising a Kozak sequence, a start codon, the coding sequence of the peptide and a stop codon, flanked by the overhangs for HindIII and AgeI, into pEGFP-N1 cut with the same enzymes. For the constructs expressing peptides with a signal sequence, a modified version of the signal sequence of HLA-E (with the final codon for alanine [GCG] replaced by one for proline [CCA] to create an MscI site [TGGCCA]) was first inserted into pEGFP-N1 using HindIII and AgeI. Annealed oligos encoding alanine followed by the desired peptide and a stop codon, and with the overhang for AgeI at the 3’ end, were inserted after the signal sequence using the MscI and AgeI restriction sites. SPP expression constructs were made by inserting cDNAs amplified from 293T cells (see below) into pcDNA3.1neo (Life Technologies) using the HindIII and NotI sites included in the PCR primers. The catalytically-inactive SPP D219A mutant was created using overlap-extension PCR (25) with KOD Hot Start Polymerase (Merck) and appropriate primers. PCR products were purified using QIAquick spin columns (QIAGEN). The sequences of the oligonucleotides used are shown in Table S1. The HCV Core-E1 expression plasmid was a kind gift of Andrea Magri (University of Oxford). All plasmids were prepared using QIAprep mini spin columns (QIAGEN), and sequences were verified by Sanger sequencing using an ABI3770.

### Cell culture and transfection

HEK 293T cells were maintained between 10% and 90% confluency at 37°C/5% CO2 in DMEM (Life Technologies) supplemented with 10% foetal bovine serum (Sigma), and penicillin/streptomycin (50 units/ml and 50 µg/ml, respectively; Life Technologies). Transfections were carried out in 6 well plates with cells at 50–70% confluency using GeneJuice (Merck) with 1 μg of total plasmid DNA per well, as per the manufacturer’s instructions.

### Gene inactivation by CRISPR/Cas9

Guide RNAs for TAP1, TAP2, and HM13 (Table S2) were selected from a previously published set (26) and inserted into pspgRNA (Addgene plasmid # 47108, a gift from Charles Gersbach) (27). HEK 293T cells were co-transfected with equal amounts of the pspgRNA plasmids and pCas9_GFP (Addgene plasmid #44719, a gift from Kiran Musunuru) (28). EGFP-positive single cells were sorted 48 hours post transfection, and genomic DNA isolated using QuickExtract (Lucigen). The gRNA binding sites were amplified using KOD Hot Start Polymerase (Merck) and the primers shown in Table S3, with the following cycling conditions: 1 minute at 96°C; 5 cycles of 25 seconds at 96°C, 45 seconds at 70°C, 45 seconds at 72°C; 21 cycles of 25 seconds at 96°C, 50 seconds at 65°C, 45 seconds at 72°C; 4 cycles of 25 seconds at 96°C, 60 seconds at 55°C, 120 seconds at 72°C. PCR products were resolved on a 1.5% agarose gel and visualised with SYBRSafe (Life Technologies), purified using QIAquick spin columns (QIAGEN) and sequenced directly with both amplification primers. The mutations in traduced by Cas9 were deconvoluted manually or using TIDE (https://tide.deskgen.com/) (29). Deconvolutions were confirmed by cloning the PCR products using a ZERO BLUNT PCR cloning kit (Life Technologies) and sequencing individual clones.

### Protein extraction and Western blotting

Cell lysates were prepared by incubating cells in RIPA buffer (150mM NaCl, 5mM EDTA, 50mM TRIS-Cl pH 8.0, 1% IGEPAL CA-630, 0.5% Sodium Deoxycholate, 0.1% SDS) on ice for 20 minutes followed by centrifugation at 13000 rpm (4 °C) for 10 minutes to remove organelle debris. Equal volumes of lysate were mixed with 2x LDS Loading buffer (Life Technologies) and run on 10% Bis-Tris NuPage gels (Life Technologies) in 1x MOPS running buffer, then transferred to PVDF membrane using 2x NOVEX Transfer Buffer (Life Technologies) and a Trans-Blot SD semi-dry transfer cell (Bio-Rad). Membrane blocking and antibody incubations were done at room temperature with constant rotation, in 5% non-fat milk prepared from powder in PBS/0.1% Tween-20, and were washed three times (5 minutes each) after each antibody incubation with PBS/0.1% Tween-20. Rabbit anti-SPP (ab190253, Abcam) or mouse anti-Hepatitis C virus Core protein (clone C7-50, Life Technologies) were visualised using IRDye 800CW Donkey anti-Rabbit IgG or Goat anti-Mouse IgG (LiCor). The control antibody was DyLight 680 β-Tubulin Loading Control Monoclonal Antibody (Life Technologies). Membranes were imaged using a LICOR Odyssey X.

### HM13 Reverse-transcription PCR

1μg of total cellular RNA prepared using RNeasy mini columns (QIAGEN) was reverse transcribed using EvoScript Universal cDNA Master (ROCHE). PCR amplification using the primers shown in Table S1, and electrophoresis, purification and cloning of the PCR products was as before.

### Staining & flow cytometry

HEK293T cells were stained 24 hours post transfection in 100µl of Dulbecco’s modified PBS (DPBS, Sigma) at 4°C for 15 minutes with antibody 3D12 (BioLegend) at 5 ng/μl, washed twice with DPBS, stained with secondary antibody (allophycocyanin-crosslinked Goat-Anti-Mouse (H+L) F(ab’)2 fragment, Life Technologies) at 130 pg/μl in 100µl of DPBS for 15 minutes at 4°C, washed as before, and fixed in 100µl of Cytofix (BD Biosciences). Stained cells were acquired using a CyAn ADP Analyser (Beckman Coulter), and analysed using FlowJo 10 (BD).

## RESULTS

### An experimental system to dissect up-regulation of HLA-E by UL40 and Rh67

Endogenous cell surface expression of HLA-E is very low, so we adopted the strategy we used previously to show that Rh67 could increase surface expression of HLA-E (19). 293T cells were transiently transfected with a plasmid expressing a single chain dimer (SCD) of HLA-E*01:01 and β2-microglobulin (Figure S1A) and expression was detected using the HLA-E-specific monoclonal antibody 3D12 (the gating strategy is shown in Figure S1B). Consistent with previous reports regarding the relative expression of the two common HLA-E alleles (30), this HLA-E*01:01 SCD expressed at lower levels than the HLA-E*01:03 SCD (Figure S1C), confirming that covalently linking β2-microglobulin to HLA-E does not significantly alter expression of HLA-E. Co-transfection of plasmids expressing both UL40 and Rh67 (protein sequences shown in Figure 1A) with the HLA-E*01:01 SCD increased cell surface expression of HLA-E (Figure 1B). A more modest, but reproducible, increase in surface expression was also seen when a plasmid expressing the HLA-A*02:01 protein is co-transfected. The up-regulation of HLA-E by Rh67 was always slightly greater than that mediated by UL40, a difference that may simply reflect the fact that the Rh67 signal sequence appears to direct translation to the ER better than that of UL40 (Figure S2). In contrast, the lower up-regulation mediated by HLA-A*02 presumably stems from its VL9 peptide being processed less efficiently than the VL9 peptides of UL40 and Rh67 as the HLA-A*02 signal sequence is equivalent to the UL40 signal sequence in directing translation to the ER.

**FIGURE 1:**
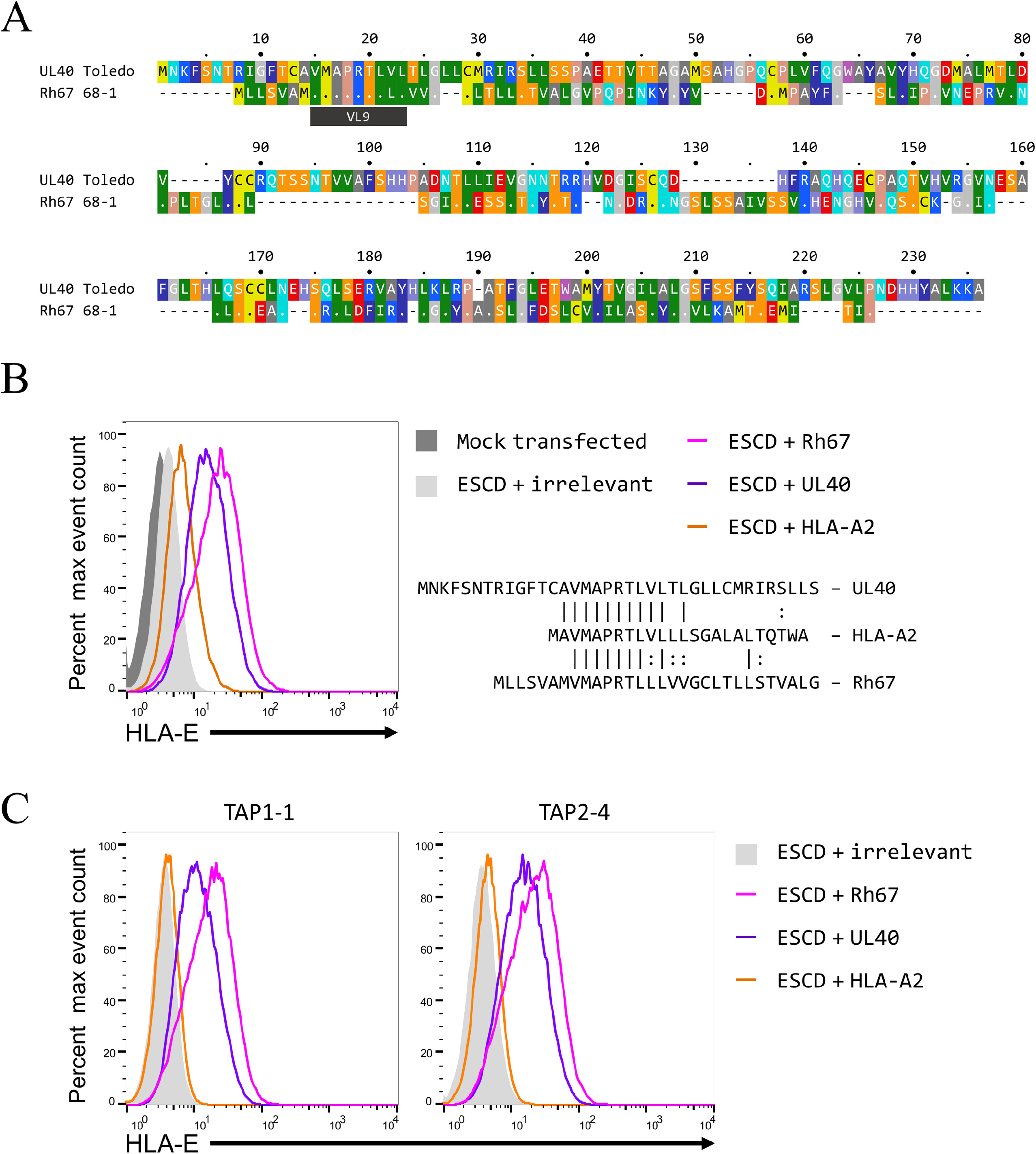
UL40 and Rh67 increase HLA-E expression by a TAP-independent pathway in 293T cells. **(A)** Alignment of the sequences of the HCMV UL40 gene (strain Toledo) and the Rhesus CMV Rh67 gene (strain 68-1). The location of the VL9 peptides, the only region of significant sequence similarity shared by the two proteins, is indicated. The sequences were aligned using EMBOSS Needle (https://www.ebi.ac.uk/Tools/psa/emboss_needle/) (46), and amino acids are coloured according to the standard RasMol scheme. **(B, C)** Representative HLA-E expression in 293T cells (panel B), or the TAP-deficient 293T clones TAP1-1 or TAP 2-4 (panel C), co-transfected with the HLA-E*01:01 SCD expression plasmid and plasmids expressing an irrelevant protein (filled light grey histogram), HLA-A*02 (orange line), UL40 (purple line), or Rh67 (pink line). Mock transfected cells are shown as the filled dark grey histogram in panel B. The sequence alignment in panel B shows a comparison of the signal sequences of the transfected UL40, Rh67 and HLA-A*02 proteins.

The UL40 mediated up-regulation of HLA-E expression has long been known not to require TAP (15, 16), and it has recently been shown that this is also true for the increase in MHC-E expression induced by Rh67 (20). To confirm that both UL40 and Rh67 were up-regulating HLA-E expression by a TAP-independent pathway in our 293T cells, we used CRISPR/Cas9 to separately inactivate the *TAP1* or *TAP2* genes (see Figure S3 and Tables S4 and S5 for details). For each, three single cell clones were established that had reduced MHC Ia expression consistent with the inactivation of TAP. Of these, clones TAP1-1 and TAP2-4 (genetic lesions shown in Figure S3E and F) were confirmed TAP deficient (Figure S3G, H and I). For both the TAP1-1 and TAP2-4 clones, we observed an almost total abrogation of the HLA-A*02-mediated up-regulation of HLA-E expression, while UL40 and Rh67 were both still able to mediate significant increases in the surface expression of HLA-E (Figure 1C). This confirms that delivery of the Rh67 peptide to HLA-E does not require transit from the cytosol to the ER lumen via TAP in 293T cells, allowing these cells to be used to dissect the sequences required for the TAP-independent increase in HLA-E expression mediated by Rh67.

### The N-terminal portion of the Rh67 signal sequence is critical for TAP-independent up-regulation of HLA-E

Up-regulation of HLA-E by UL40 is known to only require the UL40 signal sequence, and the 13 amino acids between the start codon and the UL40 VL9 peptide are both necessary for TAP-independent up-regulation of HLA-E by the UL40 VL9 peptide, and sufficient to confer TAP independence on the VL9 peptide present in the HLA-A*02 signal sequence (16) (Figure S4). As expected, replacing the Rh67 signal sequence with that of HLA-E (which does not contain the VL9 peptide) abolished up-regulation of HLA-E (Figure 2A, ESS-Rh67), whereas replacing the mature Rh67 coding sequence with that of HLA-A*02 did not (Figure 2B, RSS-A2). These results confirm that the signal sequence of Rh67 is sufficient to mediate the up-regulation of HLA-E surface expression, and that the mature protein is not required.

**FIGURE 2:**
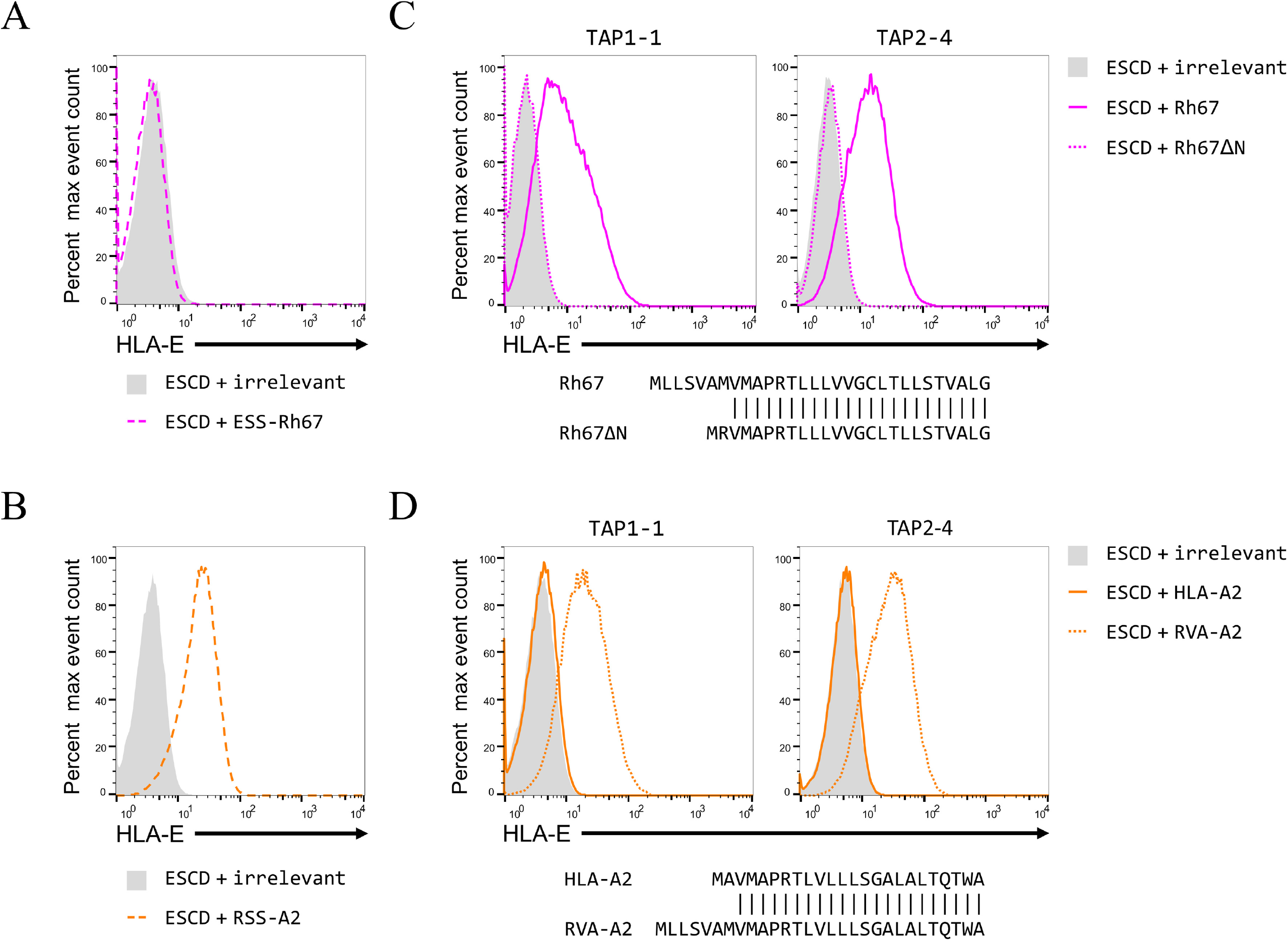
The N-terminus of the Rh67 signal sequence is critical for TAP-independent up-regulation of HLA-E. Representative HLA-E expression in 293T cells (panels **A** and **B**), or the TAP1-1 or TAP2-4 clones (panels **C** and **D**) co-transfected with the HLA-E*01:03 SCD expression plasmid and plasmids expressing an irrelevant protein (light grey filled histogram), Rh67 with the HLA-E signal sequences (ESS-Rh67; dashed pink line in panel A), HLA-A*02 with the Rh67 signal sequence (RSS-A2; dashed orange line in panel B), Rh67 (pink line in panel C), Rh67 with the N-terminal truncation shown in the sequence alignment (Rh67 ΔN; dotted pink line in panel C), HLA-A*02 (orange line in panel D), or HLA-A*02 with the first two amino acids of the signal sequence replaced with the first 7 amino acids of the Rh67 signal sequence (RVA-A2; dotted orange line in panel D).

The Rh67 signal sequence has only has 6 amino acids between its start codon and its VL9 peptide, and this sequence is less polar than the equivalent 13 amino acids in UL40. Despite this, replacing the 6 intervening amino acids in the Rh67 signal sequence with a single arginine (to preserve the expected charge distribution, as was done previously with UL40) abolished up-regulation of HLA-E by Rh67 in both the *TAP1* and *TAP2* knockout cells (Rh67ΔN, Figure 2C), without significantly affecting the functioning of the Rh67 signal sequence (Figure S2). Conversely, replacing the two amino acids upstream of the HLA-A*02 VL9 peptide with the first 7 amino acids from the N-terminus of the Rh67 signal sequence allowed TAP-independent up-regulation of HLA-E expression by the HLA-A*02 VL9 peptide (RVA-A2; Figure 2D). Thus, despite differing in both sequence composition and length from the corresponding sequence in UL40, the N-terminus of the Rh67 signal sequence is both necessary for the TAP-independence of the Rh67 VL9 peptide, and sufficient to confer TAP-independence on the HLA-A*02 VL9 peptide.

### Signal peptide peptidase is required for the processing of the UL40 and Rh67 VL9 peptides

Processing of the VL9 peptide from the MHC Ia signal sequences is known to require cleavage of the signal sequence downstream of the VL9 peptide by SPP (3). Although it was initially reported that the UL40 signal sequence was not a substrate for SPP (31), a later report suggested that it could be inefficiently processed by SPP or a SPP-like protease (16). However, it is not clear if cleavage by SPP is required to generate the UL40 VL9 peptide. Indeed, the UL40 signal sequence may have evolved to be a poor substrate if cleavage by SPP prevents proper processing of the UL40 VL9 peptide. Therefore, to address whether SPP is required for processing of the VL9 peptides in the UL40 and Rh67 signal sequences, we used CRISPR/Cas9 to knock out *HM13*, the gene that encodes SPP.

The *HM13* gRNA used targets the end of exon 4 (with the Protospacer Adjacent Motif overlapping the 5’ splice site), which encodes the longest cytoplasmic loop of SPP (Figure S5A). Four single cell clones were selected that had no SPP protein detectable by western blot (Figure S5B). Three of these clones had lesions in the *HM13* gene that would be predicted to have significant effects on expression of functional protein (Figure S5C; Table S6). The exception (clone HM13-2) had a 10 bp deletion in one allele (resulting in premature termination of translation in exon 5), and the in-frame deletion of the final 9 bp of the exon in the other. Although this clone could still express SPP with a 3 amino acid deletion, the altered sequence at the end of the exon is predicted to have a significant effect on recognition of the adjacent 5’ splice site (Table S7), likely resulting in inefficient or aberrant splicing that may well affect expression of the SPP protein.

When the *HM13* knock out clones were screened for their ability to support HLA-E up-regulation by UL40 and Rh67, two distinct phenotypes were observed (Figure 3A). None of the clones showed up-regulation of HLA-E expression by UL40, while two (HM13-1 and HM13-2) supported up-regulation by Rh67. Clones HM13-3 and HM13-4 were also unable to support maturation of the HCV Core-E1 precursor protein, which is cleaved by SPP (32), with only the immature iCore protein being detected (Figure 3B). In contrast, clone HM13-1 was indistinguishable from the parental cells, with only the mature Core (mCore) protein observed. Clone HM13-2 had an intermediate phenotype, with both mCore and iCore being detected. We conclude that clones HM13-1 and HM13-2 both retain some form of SPP activity, while clones HM13-3 and HM13-4 are truly SPP deficient.

**FIGURE 3:**
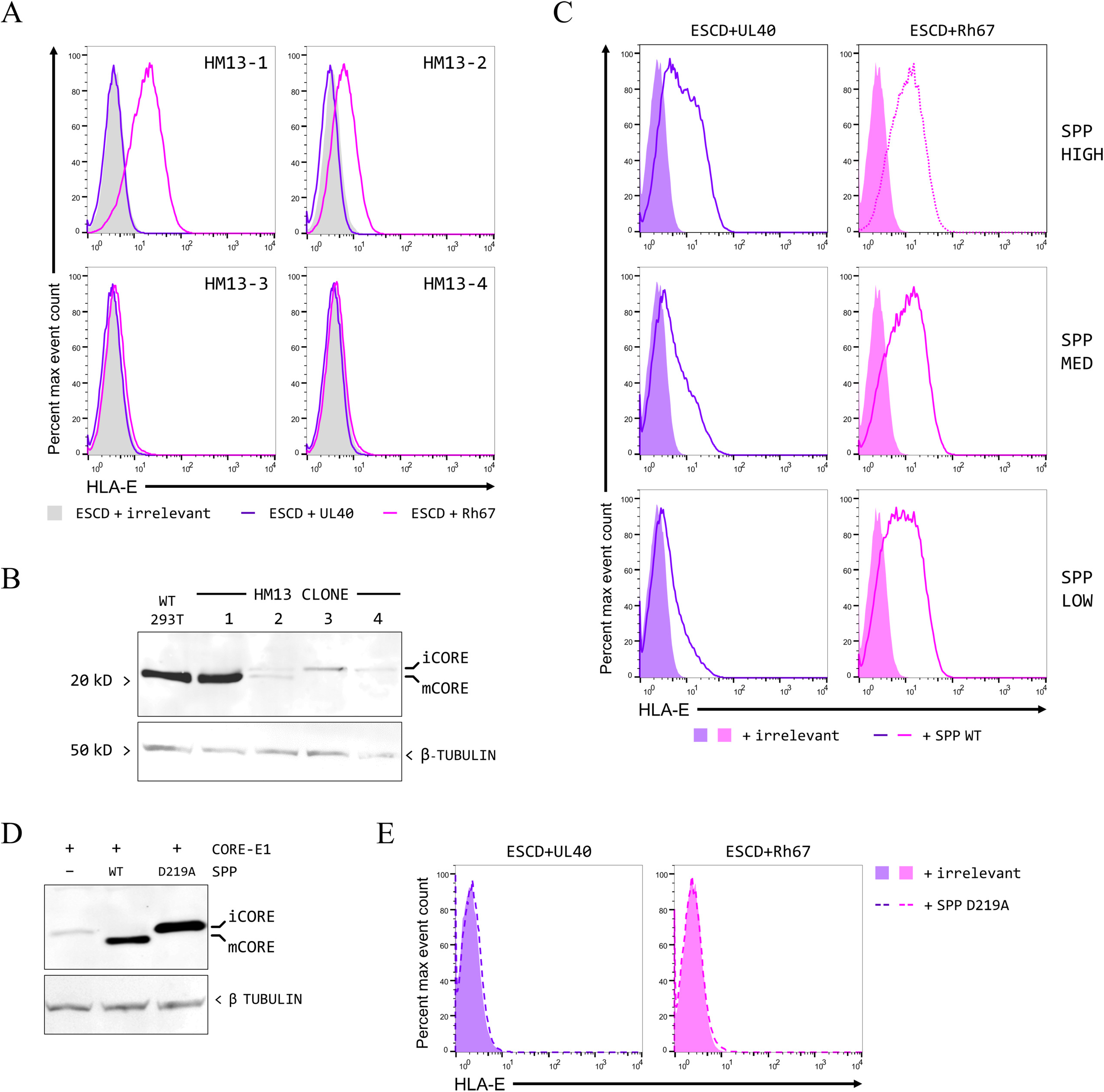
Cleavage by SPP is essential for the up-regulation of HLA-E mediated by UL40 and Rh67. **(A)** Representative HLA-E expression in the four HM13 knockout CRISPR/Cas9 single cell clones transfected with the HLA-E*01:03 SCD expression plasmid and plasmids expressing an irrelevant protein (light grey filled histogram), UL40 (purple line), or Rh67 (pink line). **(B)** Processing of the HCV Core protein, assessed by western blotting using an antibody specific for the HCV Core protein, in the parental 293T cells and all four HM13 CRISPR/Cas9 clones transfected with a plasmid expressing the Core-E1 precursor. **(C)** Representative HLA-E expression in clone HM13-3 transfected with the HLA-E*01:03 SCD expression plasmid, plasmids expressing UL40 (left-hand column, purple) or Rh67 (right-hand column, pink), as well as plasmids expressing either an irrelevant protein (filled purple or pink histograms histogram), or wild type SPP protein (purple or pink lines). Three different quantities of the wild type SPP expression plasmid were used in panel E (375ng for the top row, 187ng for the middle row, and 93ng for the bottom row, with the total amount of transfected DNA being maintained by the addition of the required quantity of an irrelevant control plasmid). **(D)** Processing of the HCV Core protein in clone HM13-3 transfected with a plasmid expressing the Core-E1 precursor, as well as plasmids expressing an irrelevant protein (denoted by “-”), wild type SPP, or a catalytically-inactive D219A SPP mutant. **(E)** Representative HLA-E expression in clone HM13-3 transfected with the HLA-E*01:03 SCD expression plasmid, plasmids expressing UL40 (left-hand column, purple) or Rh67 (right-hand column, pink), as well as plasmids expressing either an irrelevant protein (filled purple or pink histograms histogram), or the catalytically-inactive D219A SPP mutant (purple or pink lines).

The failure of UL40 and Rh67 to up-regulate expression of HLA-E in clones HM13-3 and HM13-4 suggests that processing of both the UL40 and Rh67 VL9 peptides is dependent on SPP. To confirm this, we attempted to rescue the up-regulation of HLA-E by transiently over-expressing SPP. Expression of wild type SPP in clone HM13-3 rescued up-regulation of HLA-E by both UL40 and Rh67 (Figure 3C, top row), and also cleavage of the HCV Core-E1 precursor (Figure 3D). The up-regulation of HLA-E by UL40 was more sensitive to the amount of SPP expression plasmid used in these transfections than that of Rh67 (Figure 3C, rows 2 and 3), consistent with the UL40 signal sequence being processed inefficiently by SPP. A catalytically inactive mutant of SPP (with the key aspartic acid residue at position 219 mutated to alanine) did not restore processing of the HCV Core-E1 precursor (Figure 3D), or up-regulation of HLA-E by UL40 and Rh67 (Figure 3E).

These results confirm that cleavage by SPP is essential for generating the VL9 peptides from both the UL40 and Rh67 signal sequences. Up-regulation of HLA-E expression by Rh67 in clones HM13-1 and HM13-2 could result from these cells retaining low levels of SPP activity (insufficient to support processing of UL40), or expressing SPP with altered substrate specificity as a result of the CRISPR/Cas9 lesions. It proved possible to amplify SPP cDNA from clone HM13-1, and a number of novel isoforms were identified by sequencing (Figure S6A; Table S8). The predominant isoform (HM13-1 isoform D) was almost identical to SPP isoform 1 and encoded an aberrant variant of SPP with the sequence of exon 4 (which encodes the longest cytoplasmic loop) replaced by a cryptic exon from within intron 3 (Figure S6B). However, over-expression of this isoform in clone HM13-3 rescued up-regulation of HLA-E by both UL40 and Rh67, confirming that this novel isoform could recognise both signal sequences (Figure S6C). Therefore, the impairment to UL40 mediated up-regulation in clone HM13-1 (and HM13-2) most likely stems from the low level of expression of this novel isoform, sufficient to allow processing of the VL9 peptide of Rh67, but not that of UL40.

### Improving cleavage of the UL40 signal sequence by SPP

Consistent with the UL40 signal sequence being a poor substrate for SPP, over-expression of wild type SPP in the parental 293T cells resulted in a modest increase in up-regulation of HLA-E by only UL40 (Figure 4A), suggesting that the up-regulation of surface expression by UL40 may be limited by inefficient SPP cleavage. To confirm this, we tested the effects of a previously described double mutant of the UL40 signal sequence (R31T and R33T (31) – Figure 4B) that was suggested to convert the UL40 signal sequence into an efficient SPP substrate. This mutation had little effect on the functioning of the signal sequence (Figure S2), so presumably has a direct effect on cleavage by SPP, rather than improving cleavage simply by targeting the signal sequence to the ER more efficiently. Consistent with an improvement in processing by SPP, this mutant was capable of upregulating HLA-E expression in clone HM13-1 (Figure 4C, left-hand panel). In the parental 293T cells, expression of the mutant UL40 increased up-regulation of HLA-E more than the wild type protein (Figure 5C, right-hand panel), confirming that the efficiency with which SPP cleaves the UL40 signal sequence does affect the resulting up-regulation of HLA-E.

**FIGURE 4:**
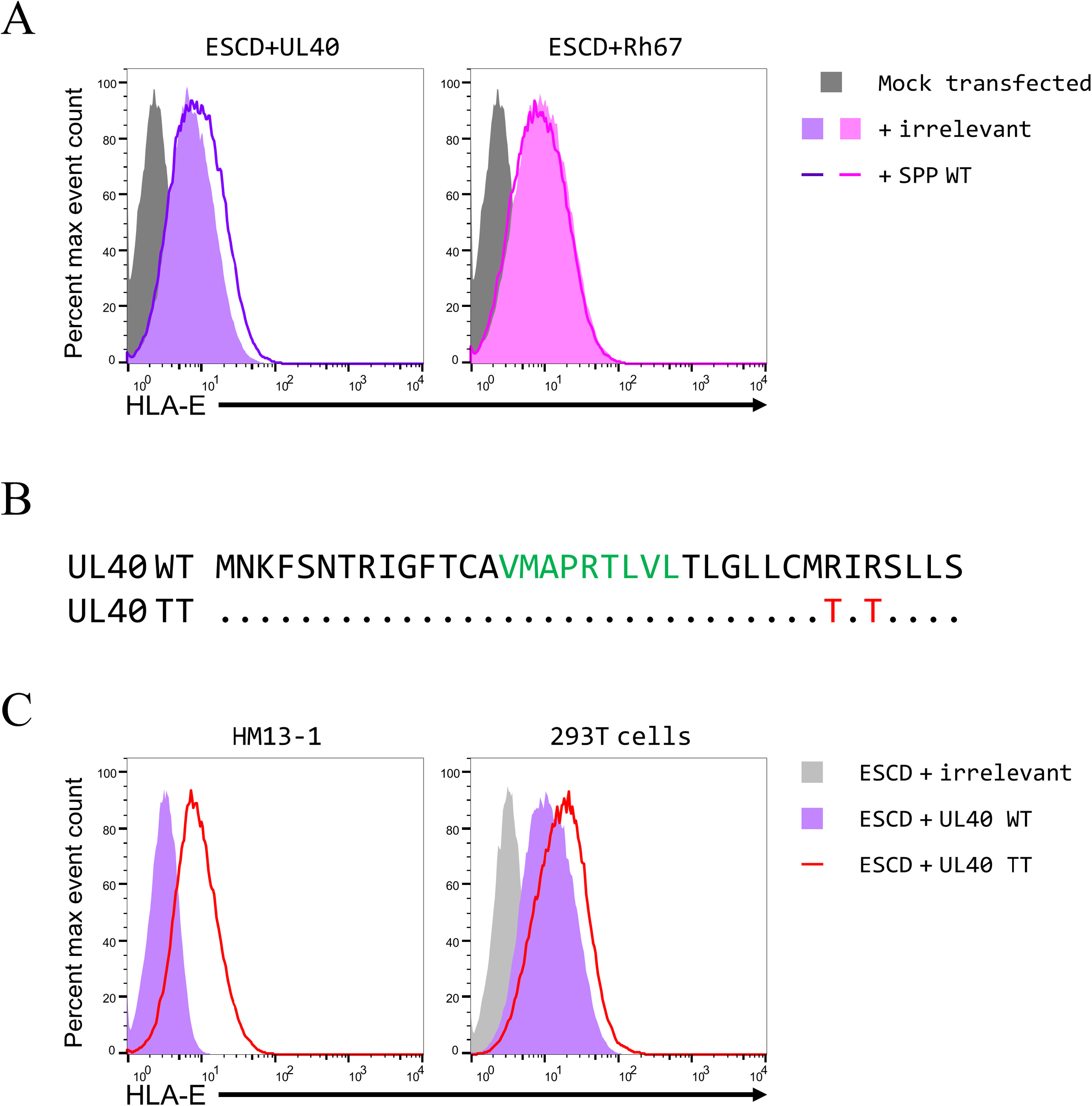
Inefficient processing of the UL40 signal sequence by SPP affects up-regulation of HLA-E. **(A)** Representative HLA-E expression in 293T cells transfected with the HLA-E*01:03 SCD expression plasmid and plasmids expressing UL40 (left-hand graph, purple) or Rh67 (right-hand graph, pink), as well as plasmids expressing an irrelevant protein (filled histogram) or wild type SPP protein (coloured line). Mock transfected cells are shown as the filled dark grey histograms. **(B)** Location of the double mutation (TT) that was previously suggested to improve cleavage of the UL40 signal sequence by SPP. **(C)** Representative HLA-E expression in clone HM13-1 (left-hand graph) or 293T cells (right-hand graph) transfected with the ESCD expression plasmid and plasmids expressing an irrelevant protein (light grey filled histograms), wild type UL40 (filled purple histogram), or the UL40 signal sequence mutant (TT, red line).

**FIGURE 5:**
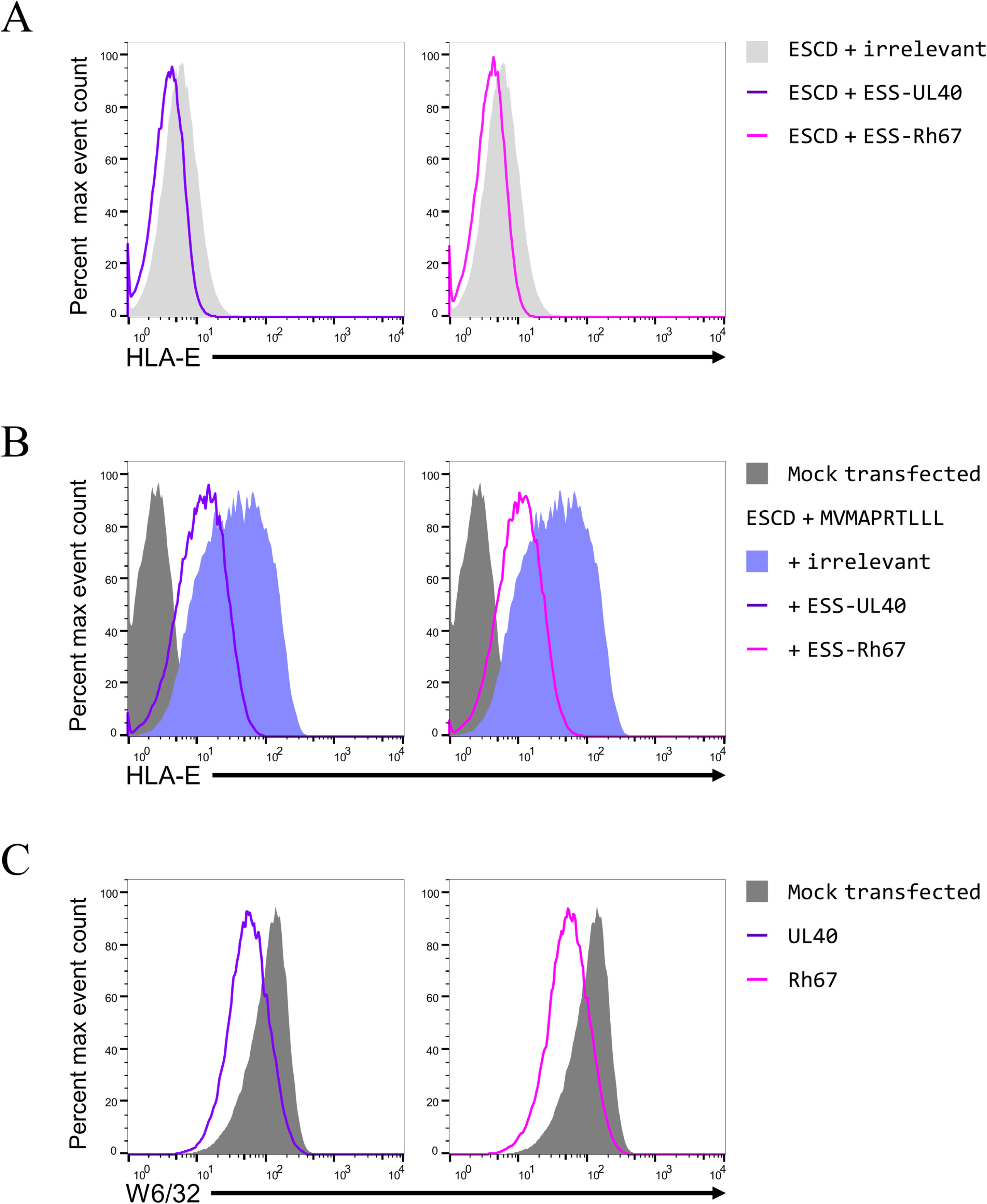
The mature UL40 and Rh67 proteins decrease surface expression of HLA-E and classical MHC class I proteins. **(A)** Representative HLA-E expression in 293T cells co-transfected with the HLA-E*01:03 SCD expression plasmid and plasmids expressing an irrelevant protein (filled light grey histogram), UL40 with the HLA-E signal sequence (ESS-UL40, purple line), or Rh67 with the HLA-E signal sequence (ESS-Rh67, pink line). **(B)** Representative HLA-E expression in 293T cells co-transfected with the HLA-E*01:01 SCD expression plasmid and a plasmid expressing VL9 peptide in the cytosol, and plasmids expressing an irrelevant protein (blue filled histogram), UL40 with the HLA-E signal sequence (ESS-UL40, purple line), or Rh67 with the HLA-E signal sequence (ESS-Rh67, pink line). **(C)** MHC class I protein expression (as assessed by W6/32 staining) of 293T cells transfected with plasmids expressing an irrelevant protein (dark grey histogram), UL40 (purple line), or Rh67 (pink line).

### The mature UL40 and Rh67 proteins reduce surface expression of MHC class I proteins

No function has been ascribed to the mature UL40 protein, and the lack of significant sequence homology means that it is not clear if the mature Rh67 protein is its functional homologue. However, in the course of these studies, we observed a slight but reproducible reduction in HLA-E expression when the HLA-E*01:03 SCD was co-expressed with UL40 or Rh67 with the HLA-E signal sequence (ESS-UL40 or ESS-Rh67, respectively – Figure 5A). A more marked reduction in surface expression of HLA-E by ESS-UL40 and ESS-Rh67 was seen when expression of the HLA-E*01:01 SCD was increased by co-expression of the VL9 peptide in the cytosol (Figure 5B). Importantly, this effect was not limited to HLA-E. Staining with the pan-MHC class I antibody W6/32 showed that expression of wild type UL40 or Rh67 (Figure 5C), or the signal sequence swapped versions (Figure S7A), resulted in a reduction in the cell surface expression of MHC Ia that was comparable to that induced by the HCMV US3 and US6 proteins (Figure S7B).

This suggests that the mature UL40 and Rh67 proteins represent previously undescribed components of the HCMV and RhCMV immune evasion arsenal, although the effect on HLA-E expression would appear to be at odds with the function of the UL40 and Rh67 signal sequences. However, the increase in HLA-E expression mediated by the natural signal sequences of UL40 and Rh67 is not dependent on TAP, in contrast to the expression of HLA-E in the experiments shown in Figure 5A and B. Therefore, we investigated if the mature UL40 and Rh67 proteins are also able to counteract the up-regulation of HLA-E expression mediated by their signal sequence VL9 peptides.

Surface expression of the HLA-E*01:01 SCD in the presence of the UL40 signal sequence (linked to HLa-A*02, USS-A2) was reduced by co-expression of either ESS-UL40 or ESS-Rh67 (Figure 6A), although HLA-E expression was still higher than when the HLA-E SCD was expressed alone. This suggests that the level of expression of HLA-E in HCMV-infected cells represents a balance between the competing processes mediated by the UL40 signal sequence and the mature UL40 protein. In contrast, the up-regulation of HLA-E expression caused by the Rh67 signal sequence (when RSS-A2 was co-expressed) was unaffected by expression of the mature UL40 or Rh67 proteins (Figure 6B). At present, it is not clear what underpins this difference in the sensitivity of the UL40 and Rh67 VL9 peptides to inhibition by the mature UL40 and Rh67 proteins, and it may suggest that expression of Mamu-E on RhCMV-infected cells may be higher than that of HLA-E on HCMV-infected cells. (2287 words)

**FIGURE 6:**
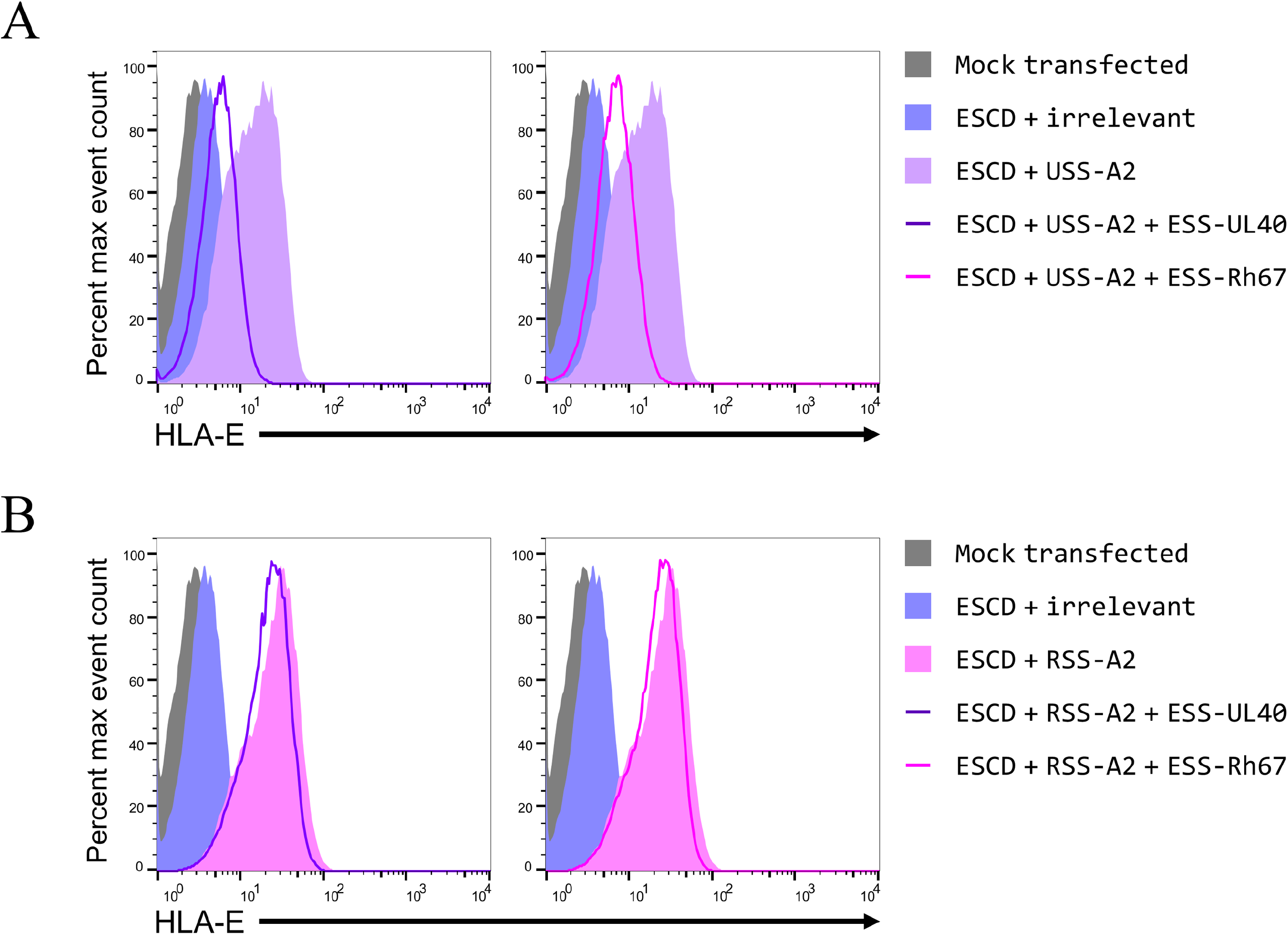
The mature UL40 and Rh67 proteins can counteract the up-regulation of HLA-E induced by the VL9 peptide of UL40 but not that of Rh67. **(A)** Representative HLA-E expression in 293T cells co-transfected with the HLA-E*01:01 SCD expression plasmid and plasmids expressing: i) only an irrelevant protein (light blue filled histogram), ii) only HLA-A*02 with the UL40 signal sequence (USS-A2, filled purple histogram), iii) HLA-A*02 with the UL40 signal sequence and UL40 with the HLA-E signal sequence (USS-A2+ESS-UL40, purple line), or iv) HLA-A*02 with the UL40 signal sequence and Rh67 with the HLA-E signal sequence (USS-A2+ESS-Rh67, pink line) **(B)** Representative HLA-E expression in 293T cells co-transfected with the HLA-E*01:01 SCD expression plasmid and plasmids expressing i) only an irrelevant protein (light blue filled histogram), ii) only HLA-A*02 with the Rh67 signal sequence (RSS-A2, filled purple histogram), iii) HLA-A*02 with the Rh67 signal sequence and UL40 with the HLA-E signal sequence (RSS-A2+ESS-UL40, purple line), or iv) HLA-A*02 with the Rh67 signal sequence and Rh67 with the HLA-E signal sequence (RSS-A2+ESS-Rh67, pink line). In both panels, mock transfected cells are shown as the filled dark grey histograms.

## DISCUSSION

We previously showed that Rh67 was likely to be the RhCMV functional homologue of the HCMV UL40 protein (19), a finding strengthened by the recent demonstration that the up-regulation of MHC-E expression mediated by Rh67 does not require TAP (20). We have now extended our characterisation of Rh67 to show striking functional similarity with UL40, despite the sequence of the two proteins showing little sequence homology. In particular, the architecture of the signal sequences appears identical: both possess N-terminal sequences that are essential for the up-regulation of HLA-E, and that confer TAP-independence on the HLA-A*02 VL9 peptide when placed at the start of the HLA-A*02 signal sequence. Moreover, despite the differences in the two signal sequences, processing of both the UL40 and Rh67 VL9 peptides also depends on cleavage by the host protease SPP. Most strikingly, however, our results also show that the mature UL40 and Rh67 proteins contribute to the broader immune evasion orchestrated by CMV by decreasing the surface expression of classical MHC class I proteins. At present, we have only studied the effects of the mature Rh67 protein on the expression of human MHC class I proteins, and it will be important to confirm that Rh67 also down-regulates Mamu-A and Mamu-B.

Unexpectedly, we also observed that the mature UL40 and Rh67 proteins can both reduce the surface expression of HLA-E. For UL40, this effect on HLA-E appears to extends to the increase in HLA-E expression mediated by its own signal sequence VL9 peptide, while the Rh67 VL9 peptide is unaffected by the action of the mature Rh67 or UL40 proteins. The UL40 signal sequence is also processed less efficiently by SPP than the Rh67 signal sequence, and this may again reduce the magnitude of HLA-E up-regulation mediated by UL40. This inefficient cleavage may simply be an indirect effect of the UL40 signal sequence being a poorer signal sequence than that of Rh67 (at least as measured by the surface expression of HLA-A*02 when its own signal sequence is replaced by those of the CMV proteins –Figure S2), resulting in a reduced amount of UL40 signal sequence available for SPP to process. The mutation that improves the UL40 signal sequence as a substrate for SPP (Figure 4) has little effect on its functioning as a signal sequence (Figure S2), however, suggesting that the reduced processing by SPP does result from the UL40 signal sequence being a poor SPP substrate.

One explanation for these differences is that UL40 may have evolved to limit the expression of HLA-E on the surface of infected cells. Perhaps related to this, the most potent peptide ligand for NKG2/CD94 interaction (VMAPRTLFL, from the HLA-G signal sequence) is rarely found in the UL40 signal sequence (33), in contrast to the other MHC Ia VL9 variants (such as VMPARTLLL, VMAPRTLIL, VMAPRTLVL, and VMAPRTVLL). Limiting HLA-E expression may be necessary to prevent recognition by NK cells expressing the activating NKG2C/CD94 ligand, which has a lower affinity for HLA-E than the inhibitory NKG2A/CD94 complex (34). The fact that Rh67 does not appear to limit its own VL9 suggests that Mamu-E expression by RhCMV-infected cells may be higher than HLA-E expression by HCMV-infected cells, perhaps reflecting differences in immune surveillance between human and macaque NK cells and the expression levels required to trigger CD94/NKG2 signalling.

At present it is not clear why the mature UL40 and Rh67 proteins can counteract the VL9 peptide of UL40 but not that of Rh67, nor how both peptides reach the lumen of the ER in the absence of TAP. The TAP-dependence of the MHC Ia VL9 peptides results from SPP cleaving the signal sequence downstream of the VL9 peptide, releasing an extended version of the peptide back into the cytosol (3). The simplest mechanism for the TAP-independence of the UL40 and Rh67 VL9 peptides, therefore, would be for SPP to cleave upstream of VL9, releasing the precursor directly into the lumen of the ER. Prediction of SPP cleavage sites is complicated by the lack of a consensus sequence, although it is likely that helix-breaking or polar residues following a stretch of alpha helix are critical (35). While the N-terminus of the Rh67 is sufficiently hydrophobic to tolerate being embedded in the ER membrane, as required for cleavage by SPP, that of UL40 is not (containing more polar residues). It was previously proposed that the N-terminus of the UL40 signal sequence would initially project into the cytosol, undergoing a topological change to allow cleavage by SPP (16). No direct evidence in support of this model was presented, and it required the UL40 signal sequence adopting a type I orientation, while SPP cleaves proteins in the type II orientation (36).

Sequences similar to that thought to be cleaved by SPP in the HLA-A*02 signal sequence (LLSG, immediately following the VL9 peptide) lie both upstream (LLSV) and downstream (LLCT) of the Rh67 VL9 peptide, although the first of these is present immediately after the start codon and may not be suitably positioned for cleavage by SPP. For UL40, the only match in the signal sequence (LLCM) lies downstream of the VL9 peptide. Therefore, for both signal sequences, it seems likely that SPP cleaves downstream of the VL9 peptide, which would release the VL9 precursors back into the cytosol. The first 5 amino acids of the Rh67 signal sequence are identical to those of a known TAP-independent epitope (<u>MLLVS</u>PLLL) which corresponds to the first 10 amino acids of the calreticulin signal sequence (37). This signal sequence can be cleaved by SPP (31), which would release a fragment containing this epitope into the cytosol. It has been proposed this peptide can enter the ER in the absence of TAP by virtue of its extremely hydrophobic sequence (16). The fragment generated by SPP cleaving after the Rh67 VL9 peptide would also be extremely hydrophobic (perhaps MLLSVAMVMAPRTLLLVV) and may behave in the same way as the calreticulin peptide. In contrast, the longer peptide that would be generated by cleavage of the UL40 signal sequence after the VL9 peptide would be significantly less hydrophobic, and presumably enters the lumen of the ER by a different mechanism, explaining the sensitivity of the UL40 VL9 peptide to the inhibitory effect of the mature UL40 and Rh67 proteins.

Having shown a requirement for SPP cleavage, we are also interested in determining what other host factors are required to process the mature VL9 peptide from both UL40 and Rh67. If precursors of these peptides do return to the cytosol, they may require processing at the C-terminus by the proteasome, as has been observed for the VL9 peptides from classical MHC class I alleles (5). Removal of the amino acids upstream of the Rh67 and UL40 VL9 peptides is also presumably required, which may suggest the involvement of the ER-resident aminopeptidases ERAP1 or ERAP2. Formation of the MHC Ia VL9 peptide also requires removal of two amino acids from the N-terminus of the signal sequence, but the enzyme responsible has yet to be identified. Disruption of ERAAP (the homologue of ERAP1) in mice has been shown to alter the peptide presented by Qa-1, the murine homologue of HLA-E (38). Rather than presenting the Qdm epitope (AMAPRTLLL) from the murine class leader sequence, ERAAP-deficient cells present a peptide derived from the Fam49b protein (FYAEATPML). However, it is not clear if this results from the absence of ERAAP preventing correct processing of the Qdm peptide, or the Fam49b peptide no longer being destroyed. Two different ERAP1 inhibitors have been shown to have no effect on surface expression of HLA-E (5), suggesting that ERAP1 may not be responsible for trimming the class I VL9 peptide in humans. A role for ERAP1 may be unlikely for the UL40 VL9 peptide, however, as a microRNA encoded in the HCMV US4 gene has been shown to downregulate expression of one of the two known isoforms of ERAP1 (39), although no equivalent microRNA has been described for RhCMV.

The level of sequence conservation between the mature UL40 and Rh67 proteins is similar to that of the other HCMV/RhCMV immunomodulatory proteins (US2/Rh182, US3/Rh184, US6/Rh185, US8/Rh186, US10/Rh187, and US11.Rh189), which show 21–30% sequence identity and 33–43% sequence similarity (17). Both proteins are known to localise in the ER (20), but it is not clear if they are associated with the membrane or present in the lumen. The TMHMM webserver (40) predicts a single transmembrane domain for Rh67, but not for UL40, and homology scanning using InterProScan (41) fails to identify any known functional domains. The structures of these proteins predicted by AlphaFold2 using the ColabFold server (42) also differ (Figure S8). The 5 models generated for UL40 show little similarity to each other when aligned, whereas those for Rh67 are much more homogeneous, differing primarily in the orientation of the C-terminus, which is predicted to form an extended alpha helix in all 5 models. Therefore, determining if these proteins are indeed functional homologues will require identification of the step(s) of the antigen processing and presentation pathway that they target.

Evasion of NK cells by the maintenance of MHC-E expression during CMV infection appears to be common through-out primate evolution, and VL9 peptides are found in all but one of the known non-human primate (NHP) CMV genomes (Table S8). As with UL40 and Rh67, the viral proteins that contain the peptide show very little sequence homology (Figure S9). In each case, the peptide is predicted to be present in a signal sequence, and its position reflects that of Rh67, with only the HCMV UL40 protein having an extended N-terminus. It seems likely that these NHP proteins will also mediate TAP-independent up-regulation of MHC-E expression, and the associated mature proteins may also play a role in decreasing surface expression of MHC class I proteins. The exception to this is the peptide present in chimpanzee CMV (TMAPKTLLL), which differs from that present in all the other NHP CMVs at two positions. The position 1 threonine is unlikely to be significant for binding by MHC-E (Figure S10), although it could affect processing of the peptide from the signal sequence, but the lysine at position 5 is striking. Interaction of HLA-E with CD94/NKG2A is abrogated by mutating position 5 of the VL9 peptide from arginine to lysine (43). Chimpanzees are the closest living relatives of humans, and their CD94 and NKG2A genes show >96% sequence identity to their human counterparts (44), and all Patr-A alleles have the canonical VL9 peptide in their leader sequences. This suggests that monitoring of class I expression by NK cells via MHC-E/CD94/NKG2A in chimpanzees will mirror that of humans. The chimpanzee CMV UL40 protein is also the most closely related to HCMV UL40 of all the NHP orthologues, but the sequence upstream of the TL9 peptide in chimpanzee CMV is not as long as that of HCMV, and not as hydrophobic as that in the other NHP CMVs. Given the extensive in vitro culturing of this stain prior to sequencing (45), it will be interesting to determine if this sequence is truly representative of chimpanzee CMV UL40, or if it reflects differences in how chimpanzee CMV evades detection by NK cells.

In conclusion, we show that despite the similarities (including the location of the critical sequences and processing of their VL9 peptides being dependent on cleavage by SPP) in how the signal sequences of UL40 and Rh67 up-regulate HLA-E expression, there are important differences. Processing of the UL40 VL9 peptide is less efficient than that of Rh67, and the resulting increase in HLA-E expression appears to be partially counteracted by the ability of the mature UL40 protein to inhibit surface expression of MHC class I. The mature Rh67 protein is also able to inhibit MHC class I expression, but does not affect the up-regulation of HLA-E mediated by its own signal sequence VL9 peptide. Such differences may have important implications for the ability of a HCMV-vectored vaccine to induce protective HLA-E-restricted responses (analogous to the protective Mamu-E responses elicited by the RvCMV 68.1 SIV vaccine) and warrants further investigation.

## ACKNOWLEDGEMENTS

We thank Andrea Margi (NDMRB, University of Oxford) for the kind gift of the HCV Core-E1 expression plasmid. Andrew Worth of the Jenner Institute for cell sorting, and Tim Rostron and John Frankland of the Weatherall Institute of Molecular Medicine Sequencing Facility for plasmid sequencing. We also thank Geraldine Gillespie and past and present members of the Gillespie, Borrow and McMichael laboratories for helpful discussions while this work was being undertaken. The work was supported by funding from the Bill and Melinda Gates Foundation (OPP1133649) and the Chinese Academy of Medical Sciences (CAMS) Innovation Fund for Medical Sciences (CIFMS 2018-I2M-2-002). P.B. and A.J.M. are Jenner Institute Investigators.

S.B. conceived and performed the experiments and analysed the data. P.B. and A.J.M. supervised the work and analysed data. S.B. wrote the manuscript, which was reviewed, edited and approved by all authors. The authors declare that they have no conflicts of interest with respect to the publication of this manuscript.

