## Supplementary Figures and Tables for "Regulation of the cell surface expression of classical and non-classical MHC proteins by the human cytomegalovirus UL40 and rhesus cytomegalovirus rh67 proteins"

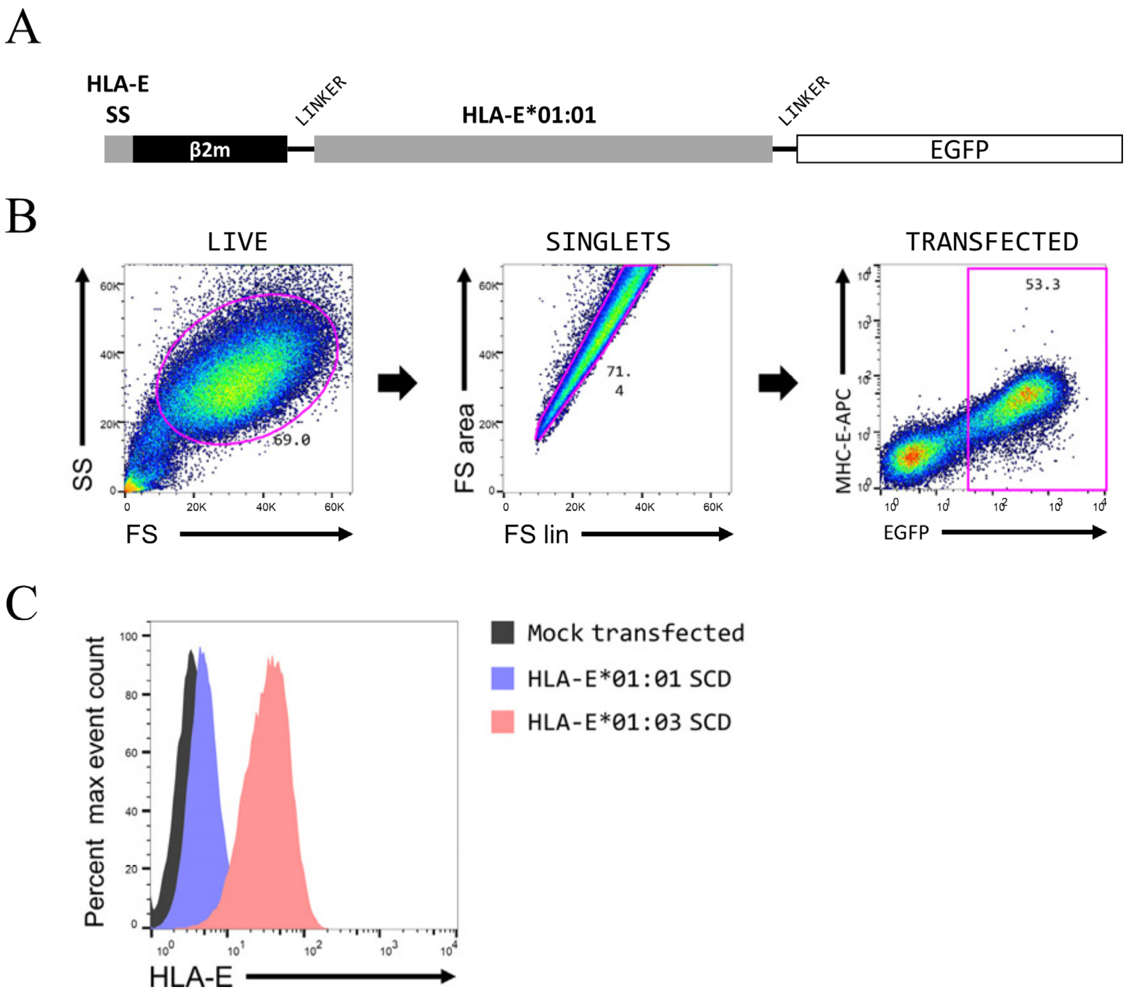


***FIGURE S1: Assaying up-regulation of HLA-E expressed as single chain dimers with β2-microglobulin***

**(A)** Schematic of the HLA-E single chain dimers (SCD) used, which comprise the HLA-E signal sequence (SS), the mature coding sequence of β2-microglobulin, a flexible Glycine-Serine linker ($[GGGGS]_4$), the mature coding sequence of HLA-E*01:01 (arginine at position 107) or HLA-E*01:03 (glycine at position 107), a second short linker (DPVAT), and EGFP. **(B)** Gating of 293T cells transfected with the HLA-E*01:03 SCD: debris is excluded on the basis of forward scatter and side scatter (FS and SS, respectively), singlets identified by FS (area) against FS (linear), and transfected cells selected by expression of EGFP. **(C)** Comparison of cell surface expression of the HLA-E*01:01 and HLA-E*01:03 single chain dimers in 293T cells (filled blue and red histograms, respectively). Mock transfected cells are shown as the dark grey filled histogram.

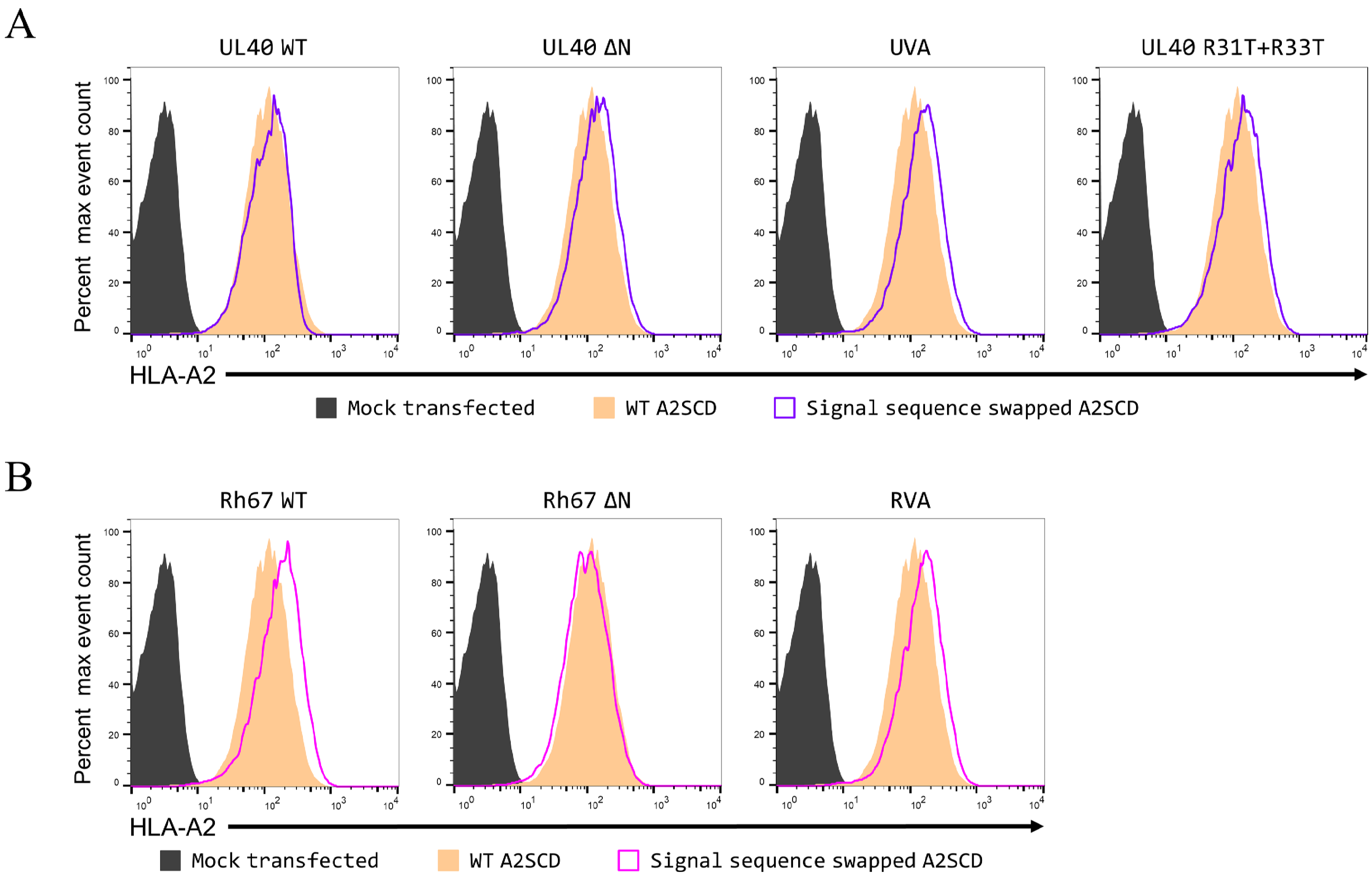
A
UL40 WT
UL40 ΔN
UVA
UL40 R31T+R33T
Percent max event count
HLA-A2
Mock transfected
WT A2SCD
Signal sequence swapped A2SCD
B
Rh67 WT
Rh67 ΔN
RVA
Percent max event count
HLA-A2
Mock transfected
WT A2SCD
Signal sequence swapped A2SCD



*FIGURE S2*

***FIGURE S2: Comparison of the HLA-A*02, UL40, Rh67, and mutated signal sequences.***

Representative staining (using the HLA-A*02-specific antibody BB7.2) of 293T cells deficient in β2-microglobulin (Brackenridge *et al.* 2022) transfected with plasmids expressing a single chain dimer of HLA-A*02 and β2-microglobulin (A2SCD) with various versions of the UL40 (panel **A**) or Rh67 (panel **B**) signal sequences. Mock transfected cells are shown by the filled dark grey histograms, expression of the A2SCD with its own signal sequence by filled orange histograms, and the SCDs with the various UL40- or Rh67-derived signal sequences as coloured outline histograms (purple and pink, respectively), with the particular mutation in each case indicated above the histogram.

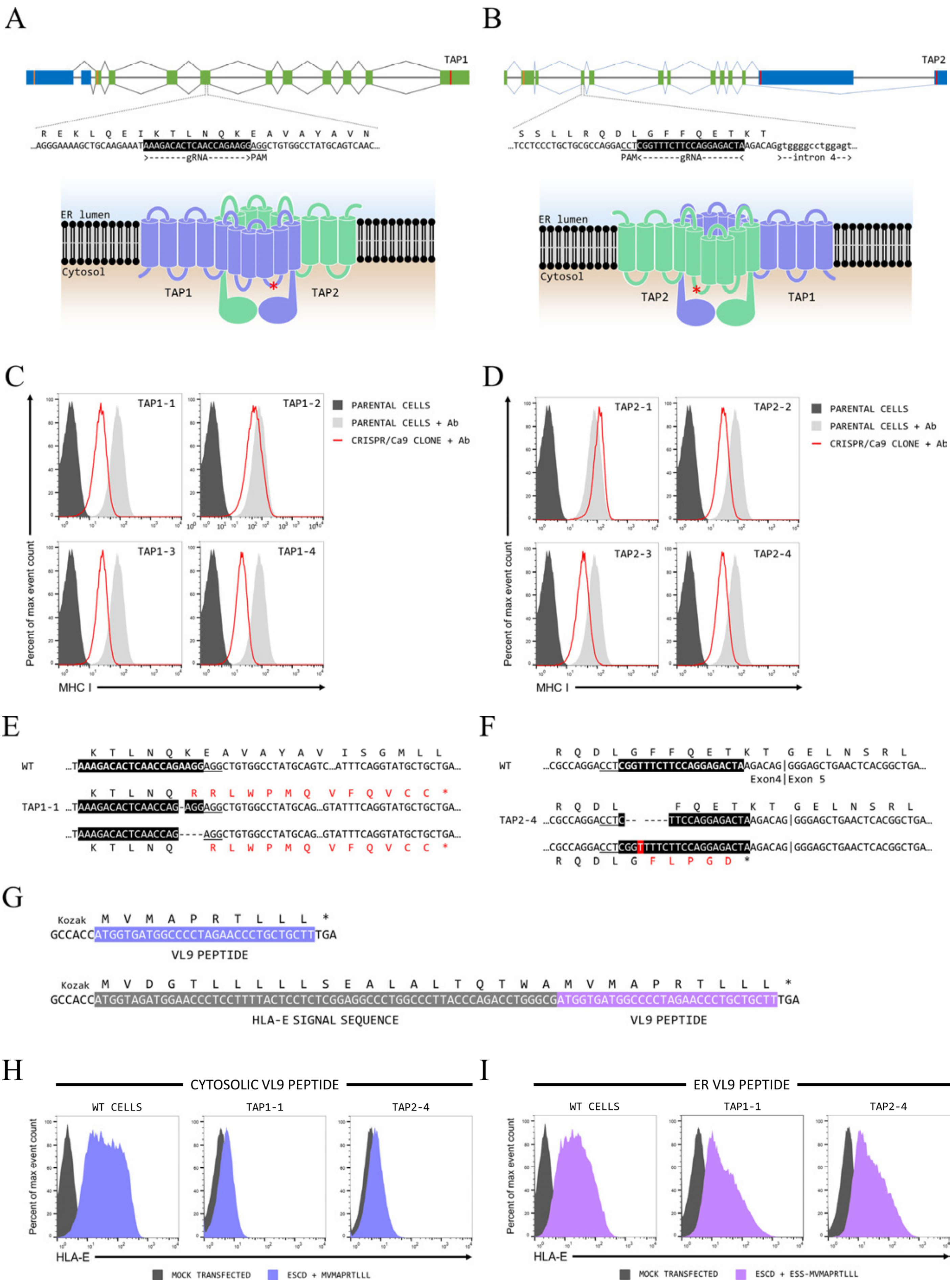
A
B
TAP1
TAP2
ER lumen
Cytosol
TAP1
TAP2
>-------gRNA------->PAM
PAM<-------gRNA-------<
>--intron 4-->
C
D
TAP1-1
TAP1-2
TAP1-3
TAP1-4
TAP2-1
TAP2-2
TAP2-3
TAP2-4
PARENTAL CELLS
PARENTAL CELLS + Ab
CRISPR/Ca9 CLONE + Ab
Percent of max event count
MHC I
E
F
WT
TAP1-1
TAP2-4
Exon4|Exon 5
G
Kozak
GCCACCATGGTGATGGCCCCTAGAACCCTGCTGCTTTGA
VL9 PEPTIDE
HLA-E SIGNAL SEQUENCE
H
I
CYTOSOLIC VL9 PEPTIDE
ER VL9 PEPTIDE
WT CELLS
HLA-E
MOCK TRANSFECTED
ESCD + MVMAPRTLLL
ESS-MVMAPRTLLL



*FIGURE S3*

***FIGURE S3: Details of the TAP1 and TAP2 CRISPR/Cas9 clones.***

**(A, B)** Schematics of the *TAP1* and *TAP2* genes and a cartoon of the TAP complex, showing the sequences and locations (red asterisk on the cartoons) of the CRISPR/Cas9 gRNA target sequences (white text on black for the sequence). **(C**, **D)** W6/32 staining of the single cell clones obtained using the *TAP1* and *TAP2* guide RNAs. Unstained single cell clones are shown as dark grey filled histograms, the stained single cell clones as the red lines, and stained parental cells as light grey filled histograms. **(E)** Sequences of the two alleles of *TAP1* in clone TAP1-1, which was selected for further study. The location of the gRNA target sequence is highlighted (white text on black), with the adjacent PAM sequence underlined. One allele has a single base deletion in the target sequence, the other a 4 base deletion starting that removes the last three bases of the target sequence and the first base of the PAM. Both lesions result in the use of the same cryptic stop codon (in the same exon as the target sequence) after the translation of 19 or 18 novel amino acids (respectively). **(F)** Sequences of the two alleles of *TAP2* in clone TAP2-4, which was selected for further study. The location of the gRNA target sequence is highlighted (white text on black), with the adjacent PAM sequence underlined. One allele of TAP2-4 has a single base insertion in the target sequence, which results in the use of a downstream cryptic stop codon in the same exon after translation of 5 novel amino acids. The other allele has a 6 base deletion starting in the target sequence, that deletes 2 amino acids from the third cytoplasmic domain of the protein, which has been implicated in peptide binding (Lehnert and Tampe 2017). Given the reduction in W6/32 staining of TAP2-4 is similar to that of clones TAP2-3 and TAP2-4, it would appear that this in-frame deletion is sufficient to abrogate TAP2 function. **(G)** The sequences encoding the peptide minigenes encoding 10mer versions of the VL9 peptide that will be expressed in the cytosol (top) or lumen of the ER (bottom). **(H, I)** Representative HLA-E expression in the parental 293T cells (left-hand graph), clone TAP1-1 (central graph) or clone TAP2-4 (right-hand graph) following transfection with the HLA-E*01:01 SCD expression plasmid a plasmid expressing either the cytosolic (panel H, filled blue histograms) or ER-targeted the MVMAPRTLLL peptides (panel I, filled purple histograms). Mock transfected cells are shown as the dark grey filled histograms.

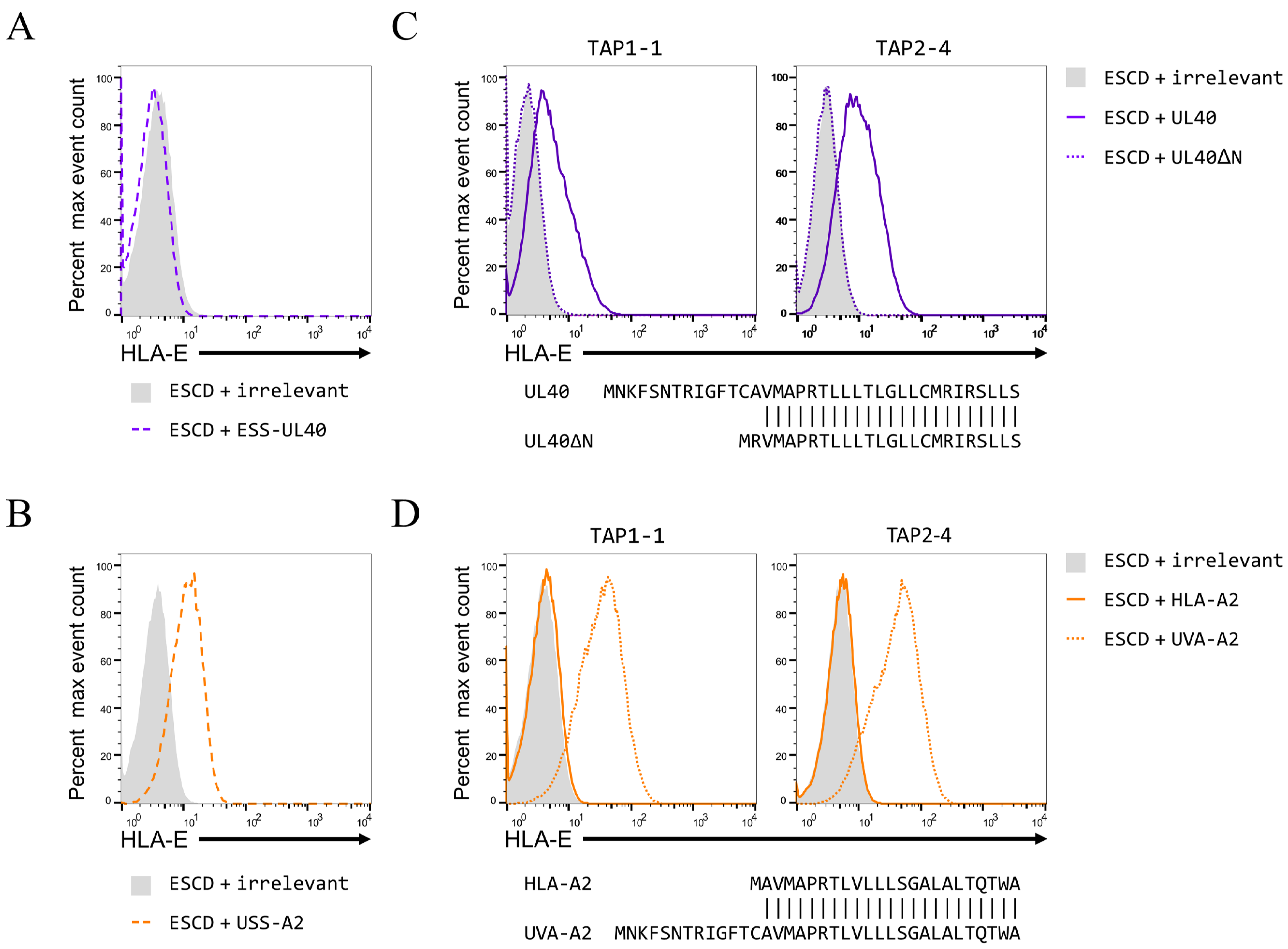
A
B
C
D
TAP1-1
TAP2-4
Percent max event count
HLA-E
ESCD + irrelevant
ESCD + ESS-UL40
ESCD + USS-A2
ESCD + UL40
ESCD + UL40ΔN
ESCD + HLA-A2
ESCD + UVA-A2
UL40 MNKFSNTRIGFTCAVMAPRTLLLTLGLLCMRIRSLLS
UL40ΔN MRVMAPRTLLLTLGLLCMRIRSLLS
HLA-A2 MAVMAPRTLVLLLSGALALTQTWA
UVA-A2 MNKFSNTRIGFTCAVMAPRTLVLLLSGALALTQTWA

*FIGURE S4*

***FIGURE S4: The N terminus of the UL40 signal sequence is critical for TAP-independent up-regulation of HLA-E***

Representative HLA-E expression in 293T cells (panels **A** and **B**), or the TAP1-1 or TAP2-4 clones (panels **C** and **D**) co-transfected with the HLA-E*01:03 SCD expression plasmid and plasmids expressing an irrelevant protein (light grey filled histogram), UL40 with the HLA-E signal sequences (ESS-UL40; dashed purple line in panel A), HLA-A*02 with the UL40 signal sequence (USS-A2; dashed orange line in panel B), UL40 (purple line in panel C), UL40 with the N-terminal truncation shown in the sequence alignment (UL40 ΔN; dotted purple line in panel C), HLA-A*02 (orange line in panel D), or HLA-A*02 with the first two amino acids of the signal sequence replaced with the first 14 amino acids of the UL40 signal sequence (UVA-A2; dotted orange line in panel D).

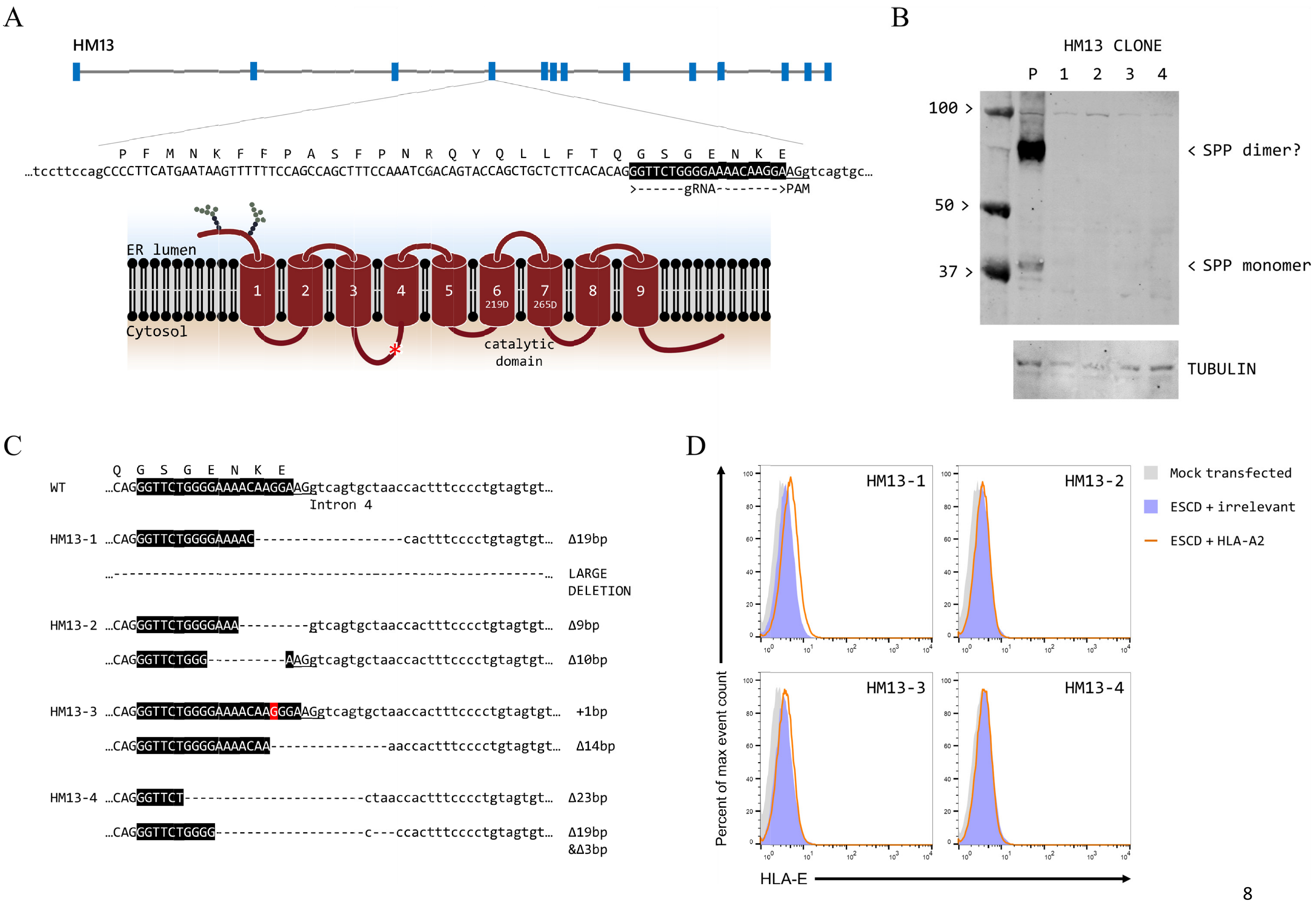
A
HM13
P F M N K F F P A S F P N R Q Y Q L L F T Q G S G E N K E
…tccttccagCCCCTTCATGAATAAGTTTTTTCCAGCCAGCTTTCCAAATCGACAGTACCAGCTGCTCTTCACACAGGGTTCTGGGGAAAACAAGGAAGgtcagtgc…
>------gRNA-------->PAM
ER lumen
Cytosol
1 2 3 4 5 6 7 8 9
219D
265D
catalytic domain
B
HM13 CLONE
P 1 2 3 4
100 >
50 >
37 >
< SPP dimer?
< SPP monomer
TUBULIN
C
Q G S G E N K E
WT …CAGGGTTCTGGGGAAAACAAGGAAGgtcagtgctaaccactttcccctgtagtgt…
Intron 4
HM13-1 …CAGGGTTCTGGGGAAAAC-------------------cactttcccctgtagtgt… Δ19bp
…------------------------------------------------------… LARGE DELETION
HM13-2 …CAGGGTTCTGGGGAAA---------gtcagtgctaaccactttcccctgtagtgt… Δ9bp
…CAGGGTTCTGGG----------AAGgtcagtgctaaccactttcccctgtagtgt… Δ10bp
HM13-3 …CAGGGTTCTGGGGAAAACAAGGGAAGgtcagtgctaaccactttcccctgtagtgt… +1bp
…CAGGGTTCTGGGGAAAACAA--------------aaccactttcccctgtagtgt… Δ14bp
HM13-4 …CAGGGTTCT-----------------------ctaaccactttcccctgtagtgt… Δ23bp
…CAGGGTTCTGGGG-------------------c---ccactttcccctgtagtgt… Δ19bp &Δ3bp
D
HM13-1
HM13-2
HM13-3
HM13-4
Mock transfected
ESCD + irrelevant
ESCD + HLA-A2
Percent of max event count
HLA-E

FIGURE S5

***FIGURE S5: Details of the HM13 CRISPR/Cas9 clones.***

**(A)** Schematic of the *HM13* gene and a cartoon of the SPP domains (with the two catalytically-important aspartic acid residues indicated), showing the sequence and location (red asterisk on the cartoon) of the CRISPR/Cas9 gRNA target sequence (white text on black for the sequence). **(B)** Western blot analysis of cell lysates of the parental cells (P) and the four HM13 CRISPR/Cas9 single cell clones. It is known that SPP forms SDS-stable dimers (Golde *et al.* 2009). **(C)** Sequences of the lesions introduced by the Cas9 protein in the two alleles of *HM13* in the four HM13 CRISPR/Cas9 single cell clones isolated. The sequence of the gene in the parental cells is shown at the top, and the location of the gRNA target sequence and PAM are indicted (white text on black and underlined, respectively). **(D)** Co-transfection of the four HM13 CRISPR/Cas9 single cell clones with plasmids expressing the HLA-E*01:01 SCD and HLA-A*02. Very slight, but reproducible, up-regulation of HLA-E surface expression was seen for clone HM13-1, but not the other three clones.

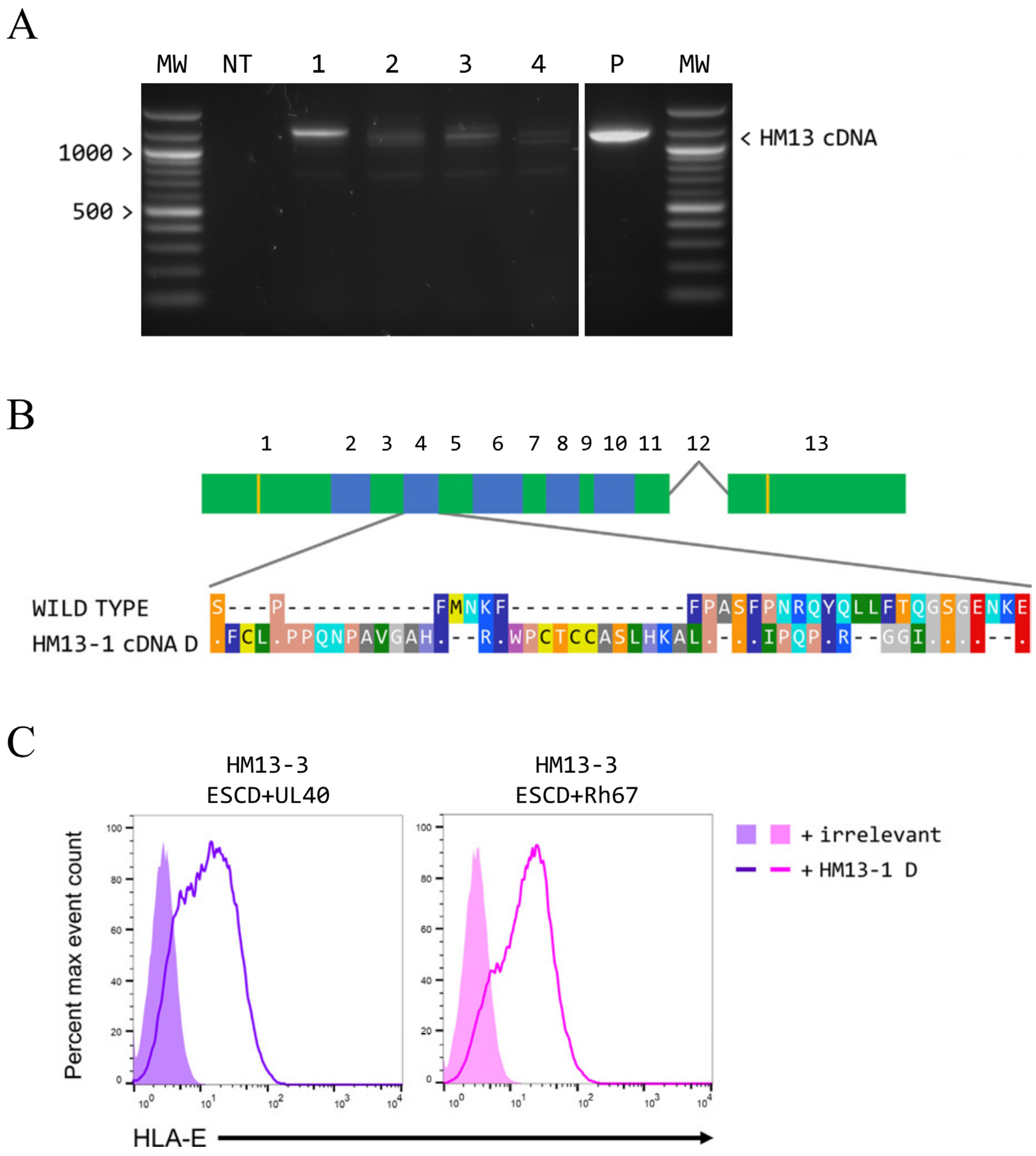


***FIGURE S6: Clone HM13-1 expresses a novel isoform of SPP that can act on both Rh67 and UL40.***

**(A)** Reverse Transcription PCR of *HM13* mRNA in the four HM13 CRISPR/Cas9 single cell clones, and the parental 293T cells (P); NT, no template control. Molecular weight (MW) markers (100 bp DNA Ladder, New England Biolabs) were run in the first and last lanes. Note that both images were from the same exposure of the same gel, but have been shown separately as intervening lanes containing irrelevant samples have been removed. **(B)** Comparison of SPP isoform 1 with the novel isoform (cDNA D) predominantly expressed by one of the *HM13* alleles in clone HM13-1. Alternating exons are shown in green and blue, with the exon in the novel allele comprising sequence from intron 3 appended to the end of exon 4. **(C)** Expression of the HLA-E in clone HM13-3 transfected with plasmids expressing the HLA-E*01:01 SCD and Rh67 (pink) or UL40 (purple), with either an irrelevant protein (filled histograms) or HM13-1 cDNA D SPP (coloured lines).

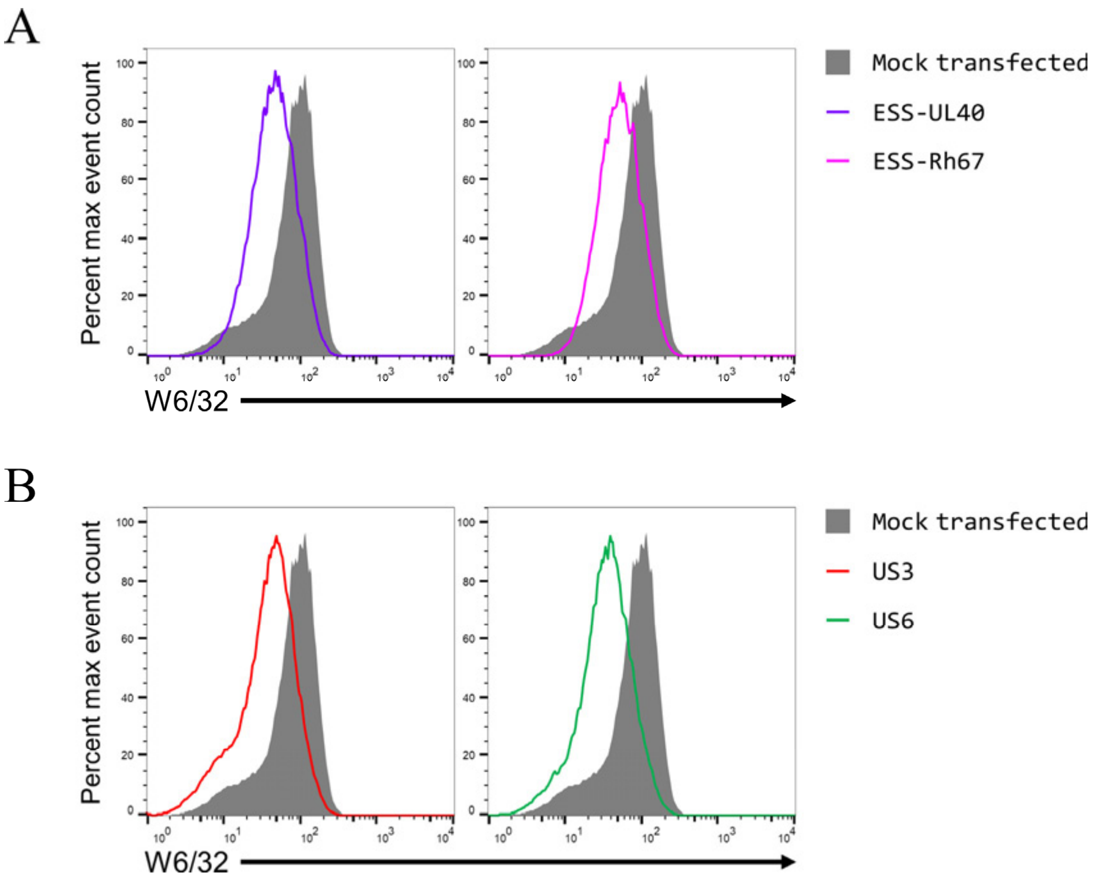


***FIGURE S7: Down-regulation of MHC class I expression by the mature UL40 and Rh67 proteins, and HCMV US3 and US6.***

Expression of MHC class I protein by 293T cells transfected with plasmids expressing ESS-UL40 or ESS-Rh67 (panel **A**, purple and pink lines, respectively), or HCMV US3 or US6 (panel **B**, red and green lines, respectively). Cells transfected with an irrelevant plasmid are shown as the filled dark grey histograms.

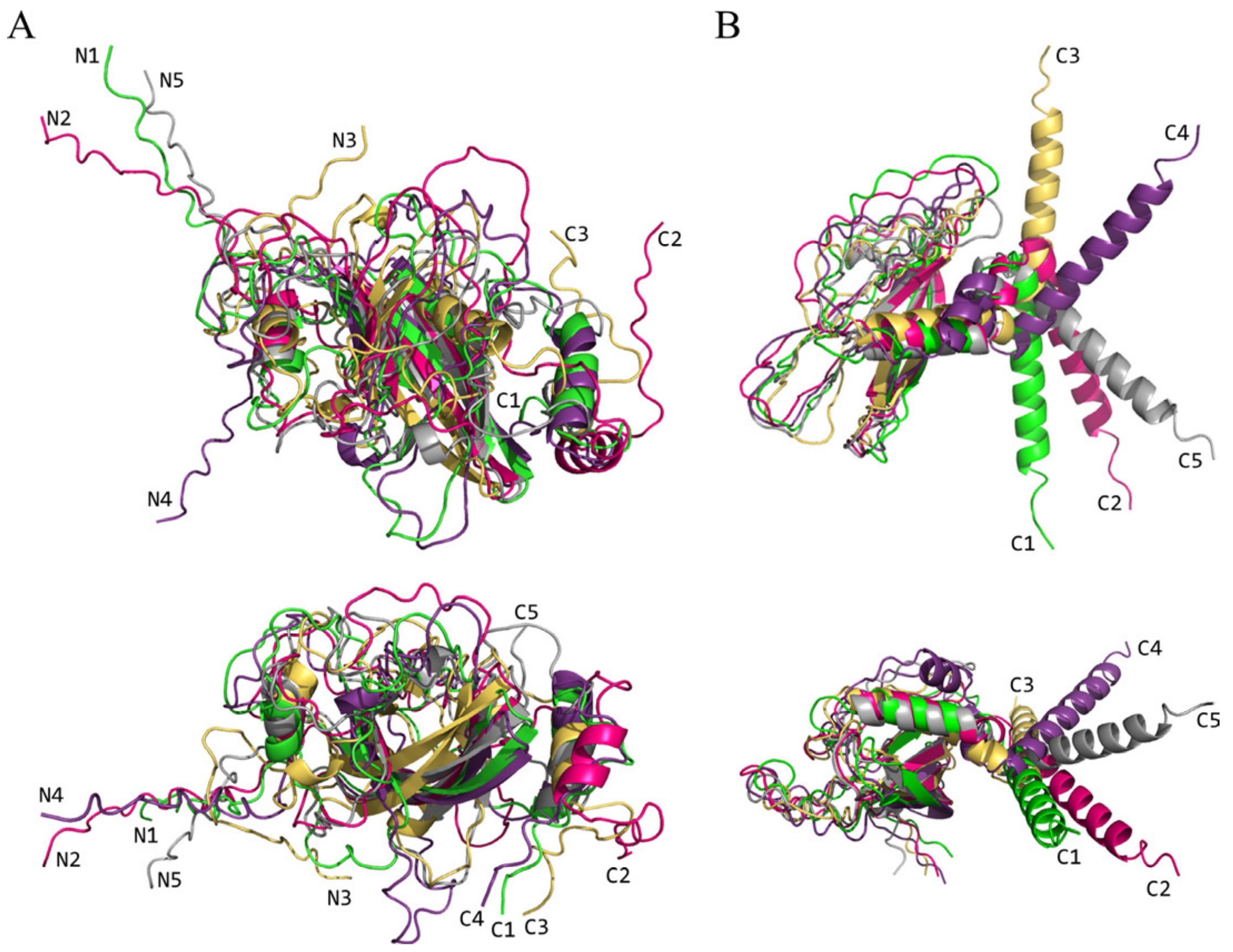


***FIGURE S8: AlphaFold2 predictions of the structures of the mature UL40 and Rh67 proteins.***

Two different views are shown of five aligned and superposed models of the structures of the mature proteins of UL40 (panel **A**) and Rh67 (panel **B**). Where possible the individual N and C termini are indicated.

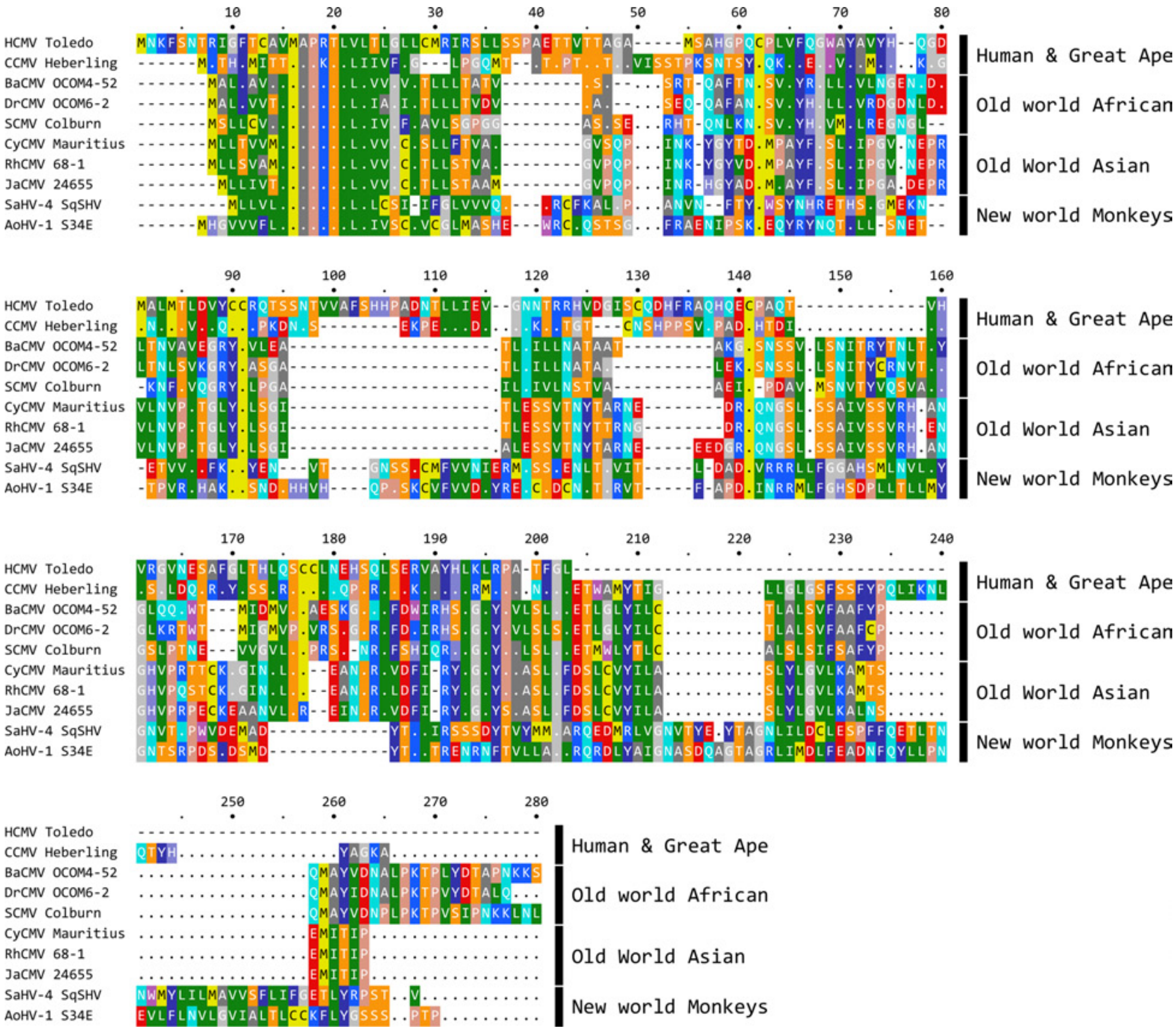


***FIGURE S9: Non-Human Primate UL40 candidates***

ClustalΩ alignment (https://www.ebi.ac.uk/Tools/msa/clustalo/) (Madeira *et al.* 2019) of the UL40 proteins encoded by 8 different NHP CMV genomes, along with HCMV UL40 (strain Toledo) and RhCMV (stain 68-1) Rh67. See Table S9 for details of the genomic sequences used.

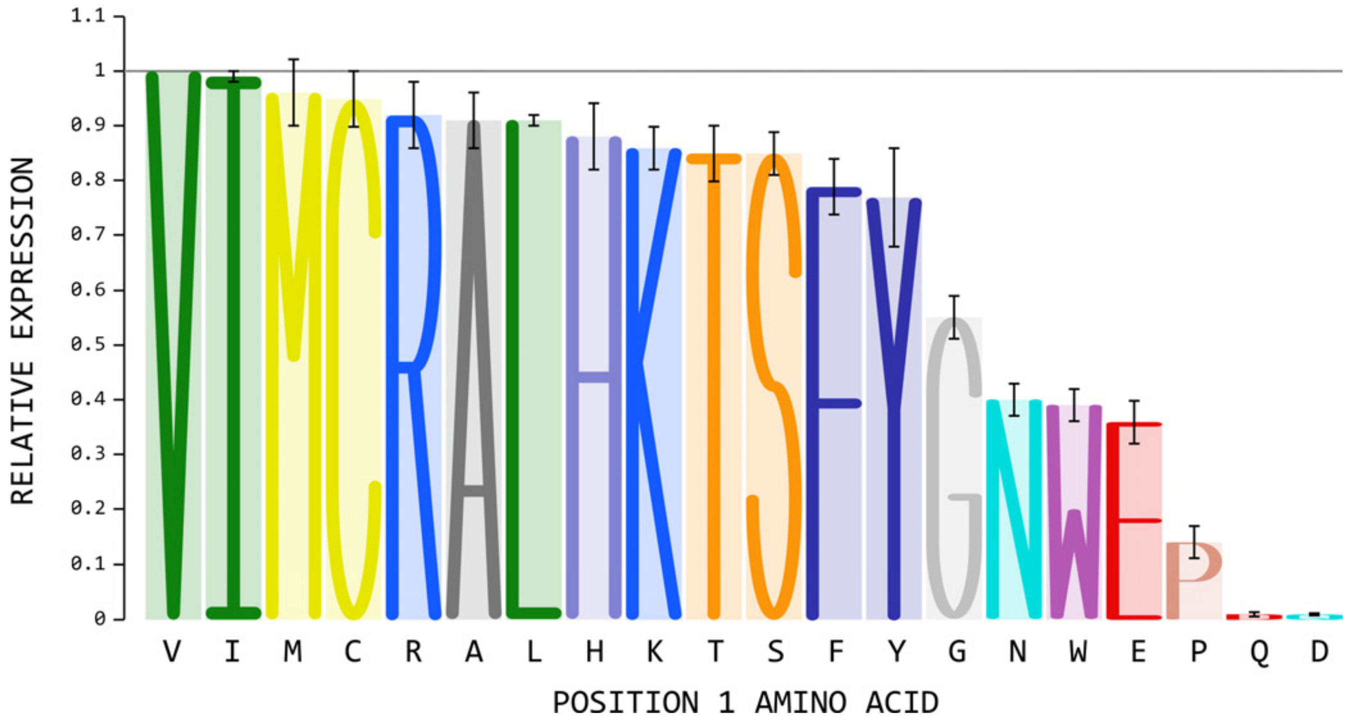


***FIGURE S10: Effects of mutating the position 1 of the VL9 peptide on expression of a single chain trimer of HLA-E*01:03, ß2-microglobulin and peptide***

Cell surface expression of single chain trimers (SCT) of HLA-E*01:03, ß2-microglobulin and the VL9 peptide (VMAPRTLLL) was used as a proxy for peptide binding affinity (Hansen et al. 2016). The first position of the peptide was changed from valine to every other amino acid, and median fluorescence intensity determined by staining with 3D12 24 hours post-transfection with the SCT expression plasmids. The results shown are the average of three independent repeats, with error bars corresponding to 4xSEM.

### TABLE S1:
### EXPRESSION CONSTRUCT PRIMERS

| CONSTRUCT | | PRIMER | SEQUENCE (5’-3') |
|---|---|---|---|
| PEPTIDE EXPRESSION | MVL9 | FORWARD | AGCTTGCCACCATGGTGATGGCCCCTAGAACCCTGCTGCTTTGAA |
| | | REVERSE | CCGGTTCAAAGCAGCAGGGTTCTAGGGGCCATCACCATGGTGGCA |
| | ESS-MVL9 | FORWARD | GCAATGGTGATGGCCCCTAGAACCCTGCTGCTTTGAA |
| | | REVERSE | CCGGTTCAAAGCAGCAGGGTTCTAGGGGCCATCACCATTGC |
| UL40 | WT | FORWARD | TCAAGCTTGCCACCATGAACAAATTCAGCAACACTCGTATC |
| | | REVERSE | GTGGATCCGCCTTTTTCAAGGCGTAGTGATGATCG |
| | TT | FORWARD | CTCCTGTGTATGACGATCACGAGTTTATTGT |
| | | REVERSE | GACAATAAACTCGTGATCGTCATACACAGGAG |
| | ΔN | FORWARD | TCAAGCTTGCCACCATGCGCGTTATGGCTCCGCGGACTTTAGTTC-<br>TGACGCTTGGAC |
| | | REVERSE | AS WT |
| Rh67 | WT | FORWARD | TCAAGCTTGCCACCATGCTGCTCAGCGTGGCGATGGTGATGG |
| | | REVERSE | GTGGATCCGGAATGGTTATCATTTCAGAAGTCATC |
| | ΔN | FORWARD | TCAAGCTTGCCACCATGCGCGTGATGGCTCCTAGGACTCTGCTTC-<br>TGGTGGTGG |
| | | REVERSE | AS WT |
| HLA-A2 | UVA-A2 | FORWARD | TCAAGCTTGCCACCATGAACAAATTCAGCAACACTCGTATCGGCT-<br>TCACTTGCGC-GGTCATGGCGCCCCGAACCCTCGTCC |
| | | REVERSE | GTGGATCCACTTTACAAGCTGTGAGAGA |
| | RVA-A2 | FORWARD | TCAAGCTTGCCACCATGCTGCTCAGCGTGGCGATGGTCATGGCGC-<br>CCCGAACCCTCGTCC |
| | | REVERSE | AS UVA-A2 |
| HM13 (SPP) | WT | FORWARD | TTAAGCTTGCCACCATGGACTCGGCCCTCAGCGATCCGCA |
| | | REVERSE | GAGCGGCCGCTCATTTCTCTTTCTTCTCCAGCCCCTTCG |
| | D219A | FORWARD | CTCTTCATCTACGCTGTCTTCTGGGTATTTGG |
| | | REVERSE | CAAATACCCAGAAGACAGCGTAGATGAAGAG |

**NOTE**: The locations of mutations and restriction sites (full or partial) used for cloning are underlined

### TABLE S2:
### CRISPR/CAS9 GUIDE RNA TARGETS

| GENE | SEQUENCE (5'-3') |
|---|---|
| *TAP1* | GAAAGACACTCAACCAGAAGG |
| *TAP2* | GTAGTCTCCTGGAAGAAACCG |
| *HM13* | GGTTCTGGGGAAAACAAGGA |

**NOTE:** Additional guanosines were added to the start of the *TAP1* and *TAP2* gRNAs to ensure efficient transcription

### TABLE S3:
### GENOMIC PCR PRIMERS USED

| GENE | FORWARD PRIMER (5'-3') | REVERSE PRIMER (5'-3') |
|---|---|---|
| *TAP1* | TGGTCCATGTTCCCAGGTTGCTC | CCTGAGAGGCAAAGGAAGGCCC |
| *TAP2* | CGCTGGGACAGAAGCAAGCA | ACAGCCCCACCACTTTCACCA |
| *HM13* | GCTCTGTTTGCCGACTTGCT | CCCAGGATCTCCACACTTCCC |

### TABLE S4:
### GENETIC LESIONS OBSERVED IN THE ALLELES OF THE *TAP1* CRISPR/CAS9 SINGLE CELL CLONES

| CLONE | ALLELE 1 | ALLELE 2 |
|---|---|---|
| TAP1-1 | Δ1bp<br>Premature stop in exon 6 | Δ4bp<br>Premature stop in exon 6 |
| TAP1-2 | Δ3bp:<br>1 amino acid deleted | Δ4bp<br>Premature stop in exon 6 |
| TAP1-3 | Δ2bp<br>Premature stop in exon 5 | Δ15bp<br>5 amino acids deleted |
| TAP1-4 | +1bp<br>Premature stop in exon 5 | Δ11bp<br>Premature stop in exon 5 |

### TABLE S5:
### GENETIC LESIONS OBSERVED IN THE ALLELES OF THE *TAP2* CRISPR/CAS9 SINGLE CELL CLONES

| CLONE | ALLELE 1 | ALLELE 2 |
|---|---|---|
| TAP2-1 | UNMUTATED | +1bp<br>premature stop in exon 4 |
| TAP2-2 | +1bp<br>Premature stop in exon 4 | +6bp<br>2 amino acids added |
| TAP2-3 | Δ6bp<br>2 amino acids deleted | Δ6bp<br>2 amino acids deleted |
| TAP2-4 | +1bp<br>Premature stop in exon 4 | Δ6bp<br>2 amino acids deleted |

### TABLE S6:
### GENETIC LESIONS OBSERVED IN THE ALLELES OF THE *HM13* CRISPR/CAS9 SINGLE CELL CLONES

| CLONE | ALLELE 1 | ALLELE 2 |
|---|---|---|
| HM13-1 | Δ19bp<br>exon 4 splice site removed | LARGE DELETION[1] |
| HM13-2 | Δ9bp<br>3 amino acids deleted[2] | Δ10bp<br>Premature stop in exon 5 |
| HM13-3 | +1bp<br>premature stop in exon 5 | Δ14bp<br>Exon 4 splice site removed |
| HM13-4 | Δ19bp<br>exon 4 splice site removed | Δ23bp & Δ3bp<br>Exon 4 splice site removed |

**NOTES:**

1. The full extent of the deletion has not been determined, but it extends upstream of the target sequence into intron 3.
2. The three amino acids deleted are not contiguous: …GSGENKEEII… becomes …GSGEKII...

### TABLE S7:
### 5' SPLICE SITE PREDICTION FOR EXON 4 OF THE *HM13* GENE IN THE PARENTAL CELLS AND CLONE HM13-2

| SPLICE SITE | SEQUENCE | MaxENT SCAN[1] | BDGP SP[2] |
|---|---|---|---|
| Optimal 5' splice site | …CAGgtaagt… | 10.86 | N/A[3] |
| Exon 4 *HM13* WT | …AAGgtcagt… | 8.68 | 0.93 |
| Exon 4 *HM13* clone 2 | …AAAgtcagt… | 1.98 | NOT PREDICTED |

**NOTES:**
1. http://hollywood.mit.edu/burgelab/maxent/Xmaxentscan_scoreseq.html
2. https://www.fruitfly.org/cgi-bin/seq_tools/splice.pl
3. N/A = not applicable (splice sites predicted in situ)

### TABLE S8:
### cDNA SPECIES IDENTIFIED IN CLONE HM13-1

| cDNA | COUNT | SEQUENCE | ALLELE | PROTEIN |
|---|---|---|---|---|
| A | 1/19 | Exon 3 extended by 98bp, exons 4–6 skipped, premature stop codon in exon 3 | 1 | Non-functional |
| B | 2/19 | Exon 4 skipped, premature stop codon in exon 5 | 1 | Non-functional |
| C | 2/19 | Exons 3–6 skipped, premature stop codon in exon 12 | 1 | Non-functional |
| D | 12/19 | Exon 4 replaced by a novel (in frame) exon comprised of 136bp from intron 3, followed by GG, then the last 5bp of exon 4. | 2 | Functional |
| E | 1/19 | As D but with exon 11 skipped, premature stop codon in exon 12 | 2 | Non-functional |
| F | 1/19 | As D but with exon 3 extended by 98bp (as in A), premature stop codon in exon 3 | 2 | Non-functional |

**TABLE S9:**

**NON-HUMAN PRIMATE CMV PROTEINS CONTAINING VL9 OR VL9-LIKE PEPTIDES**

| SPECIES | GENBANK | SEQUENCE | GENE | PEPTIDE |
|---|---|---|---|---|
| *Pan troglodytes* (Chimpanzee) | AF480884 | Panine herpesvirus 2 strain Heberling, complete genome | UL40 | TMAPKTLLI |
| *Papio ursinus* (Baboon) | MT157321 | Baboon cytomegalovirus isolate 31282, complete genome | UL40 | VMAPTRLLL |
| | MT157322 | Baboon cytomegalovirus isolate 34826, complete genome | UL40 | VMAPTRLLL |
| | KR351281 | *Papio ursinus* cytomegalovirus isolate OCOM4-52, complete genome | n/a | VMAPTRLLL |
| *Mandrillus leucophaeus* (Drill) | KR297253 | *Mandrillus leucophaeus* cytomegalovirus isolate OCOM6-2, complete genome | n/a | VMAPRTLLL |
| *Cercopithecus aethiops* (African Green Monkey) | FJ483969 | Cercopithecine herpesvirus 5 strain Colburn, complete genome | UL40 | VMAPRTLLL |
| | FJ483968 | Cercopithecine herpesvirus 5 strain 2715, complete genome | UL40 | VMAPRTLLL |
| | U27469 | Stealth virus 1 strain ATCC VR-2343 | n/a | VMAPRTLLL |
| *Macaca fascicularis* (Cynomolgus Macaque) | JN227533 | Cynomolgus macaque cytomegalovirus strain Ottawa, complete genome | Cy63 | VMAPRTLLL |
| | KP796148 | Cynomolgus macaque cytomegalovirus strain Mauritius, complete genome | Cy63 | VMAPRTLLL |
| | KX689263 | Cynomolgus cytomegalovirus isolate 31906, complete genome | Cy67 | VMAPRTLLL |
| | KX689264 | Cynomolgus cytomegalovirus isolate 31907, complete genome | Cy67 | VMAPRTLLL |
| | KX689265 | Cynomolgus cytomegalovirus isolate 31908, complete genome | Cy67 | VMAPRTLLL |
| | KX689266 | Cynomolgus cytomegalovirus isolate 31907, complete genome | Cy67 | VMAPRTLLL |
| | MT157323 | Cynomolgus cytomegalovirus strain 31709, complete genome | Cy67 | VMAPRTLLL |
| *Macaca fuscata* (Japanese Macaque) | MT157324 | Japanese cytomegalovirus strain 24655, complete genome | Ja67 | VMAPRTLLL |
| *Saimiri sciureus* (Squirrel Monkey) | FJ483967 | Saimiriine herpesvirus 4 strain SqSHV, complete genome | UL40 | VMAPRTLLL |
| *Aotus trivirgatus* (Owl Monkey) | FJ483970 | Aotine herpesvirus 1 strain S34E, complete genome | UL40 | VMAPRTLLL |

***NOTES:***
1. *n/a = not annotated in the GenBank entry*
2. *This table does not include any of the known Rhesus CMV Rh67 sequences*
3. *An alignment of representative sequences for each species is shown in Figure S9*
